# Spatially selective stimulation of the pig vagus nerve to modulate target effect versus side effect

**DOI:** 10.1101/2022.05.19.492726

**Authors:** Stephan L. Blanz, Eric D. Musselman, Megan L. Settell, Bruce E. Knudsen, Evan N. Nicolai, James K. Trevathan, Ryan S. Verner, Jason Begnaud, Aaron J. Suminski, Justin C. Williams, Andrew J. Shoffstall, Warren M. Grill, Nicole A. Pelot, Kip A. Ludwig

## Abstract

Electrical stimulation of the cervical vagus nerve using implanted electrodes (VNS) is FDA-approved for the treatment of drug-resistant epilepsy, treatment-resistant depression, and most recently, chronic ischemic stroke rehabilitation. However, VNS is critically limited by the unwanted stimulation of nearby neck muscles – a result of non-specific stimulation activating motor nerve fibers within the vagus. Prior studies suggested that precise placement of small epineural electrodes can modify VNS therapeutic effects, such as cardiac responses. However, it remains unclear if placement can alter the balance between intended effect and limiting side effect.

We used an FDA investigational device exemption approved six-contact epineural cuff to deliver VNS in pigs and quantified how epineural electrode location impacts on- and off-target VNS activation. Detailed post-mortem histology was conducted to understand how the underlying neuroanatomy impacts observed functional responses. Here we report the discovery and characterization of clear neuroanatomy-dependent differences in threshold and saturation for responses related to both effect (change in heart rate) and side effect (neck muscle contractions). The histological and electrophysiological data were used to develop and validate subject-specific computation models of VNS, creating a well-grounded quantitative framework to optimize electrode location-specific activation of nerve fibers governing intended effect versus unwanted side effect.

## Introduction

Vagus nerve stimulation (VNS) is FDA-approved to treat multiple indications and presents a significant clinical and market opportunity for future expansion (Beekwilder & Beems, 2010; Groves & Brown, 2005). VNS is currently approved for the treatment of epilepsy, depression, obesity, and stroke rehabilitation (U.S. Food & Drug Administration, 1997, 2005, 2015, 2021). Clinical trials are evaluating VNS for other indications including heart failure, diabetes, and rheumatoid arthritis (Boston Scientific Corporation, 2021; Drewes, 2021; SetPoint Medical Corporation, 2021). Unsurprisingly, given its potential in multiple therapeutic applications, there is a substantial market for VNS. In 2018, the global market for VNS was estimated at $500 million, with projected growth to nearly $1.2 billion by 2026 (Fortune Business Insights, 2019).

The vagus nerve (VN) has such a wide range of clinical applications due in part to its complex organization consisting of over 100,000 nerve fibers in humans (Hoffman & Schnitzlein, 1961). Notably, these fibers are responsible for multiple sensorimotor functions of the neck and throat, as well as autonomic functions of the cardiac system, respiratory system, and most organs within the abdomen. Sensory afferent fibers from the visceral organs ultimately influence noradrenergic, serotonergic, and cholinergic inputs to the cortex (Dorr & Debonnel, 2006; Groves & Brown, 2005; Pavlov & Tracey, 2012). Somatic motor efferent fibers run through the pharyngeal, superior laryngeal, and recurrent laryngeal branches of the VN, which innervate the pharyngeal, cricothyroid, and cricoarytenoid muscles of the neck (Câmara & Griessenauer, 2015; Rea, 2014).

Motor activation of the neck muscles is the most common side effect in VNS implant recipients (FDA, 2017). Contemporary VNS cuff electrodes wrap ~340° (dependent on nerve diameter) around the VN, and they indiscriminately stimulate the entire cross section of the VN trunk, including both sensory and motor nerve fibers. Electrical stimulation activates large-diameter motor efferent fibers at lower thresholds than smaller diameter and unmyelinated fibers (Grill, 2004; Horch & Kipke, 2017). Therefore, VNS often elicits therapy-limiting side effects (mediated by large-diameter motor nerve fibers) at lower stimulation amplitudes than required to achieve intended therapeutic effects (mediated by medium- and small-diameter fibers) (Yoo et al., 2013). Activation of large-diameter motor nerve fibers is responsible for cough, throat pain, voice alteration, and dyspnea reported in up to 66% of patients (De Ferrari et al., 2017; Handforth et al., 1998; Nicolai et al., 2020; The Vagus Nerve Stimulation Study Group, 1995). In a clinical study of VNS to treat heart failure, desired VNS evoked heart rate responses indicative of cardiac engagement were achieved in only 12% of measurements taken at the 6- and 12-month end points using stimulation parameters (i.e., signal frequency, pulse duration, and output current) similar to those used in the treatment of epilepsy and depression; off-target stimulation of the neck muscles was the principle therapy-limiting-side effect (De Ferrari et al., 2017). Although VNS is a promising therapy for treating various disorders, its efficacy can be limited by side effects induced by activation of off-target pathways.

Recent studies of VNS suggest that smaller, multi-contact electrodes may provide better selectivity for isolating desired therapeutic effects from off-target side effects (Fitchett et al., 2021; Ordelman et al., 2013; Shulgach et al., 2021). Specifically, the studies demonstrated that the magnitude of heart rate changes may be altered by changing the electrode location around the circumference of the VN. While encouraging, the studies focused only on cardiac response and did not quantify the more common side effects (i.e., motor efferent activation of neck muscles). Further, the studies were phenomenological in design and did not document how the underlying organization of nerve fibers within the VN, coined ‘vagotopy,’ related to the functional outcomes of stimulation.

In this study, we addressed this gap by quantifying the relationship between the anatomical organization of fiber types in the pig cervical vagus nerve and the functional outcomes of electrical stimulation, including recordings of evoked neural activity, activation of deep neck muscles, and changes in heart rate. The pig is the animal model most resembling the human in terms of diameter and fascicular complexity of the cervical vagus, as well as vagal innervation of deep neck muscles (Nicolai et al., 2020; Pelot et al., 2020; Settell et al., 2020). VNS was performed using the six-contact ImThera cuff electrode (LivaNova, London, UK); the ImThera electrode was awarded an FDA Investigational Device Exemption to conduct clinical studies to stimulate the hypoglossal nerve to treat obstructive sleep apnea, indicating that promising results in application of the ImThera device to the VN could be quickly translated to future human studies. We conducted histology on the VN from each animal to trace fascicles from the nodose ganglion (i.e., a location of known functional organization) to the cervical nerve trunk (i.e., where the cuff electrode was placed); we could thus identify functional groupings of fascicles within the cuff electrode that predominantly contained either motor and parasympathetic efferents fibers or sensory afferent fibers from the visceral organs. These physiological and histological data were used to implement and validate individual-specific, anatomically realistic, biophysical computational models of contact location-specific activation. These computational models provide a conceptual and mechanistic framework for quantifying the opportunities and limits of location-specific fiber activation that govern desired changes in sympathetic/parasympathetic tone versus unwanted activation of efferent motor nerves innervating deep neck muscles.

## Materials & Methods

The Institutional Animal Care and Use Committee approved all animal care and procedures at the University of Wisconsin-Madison. Six University of Wisconsin minipigs were included in the study weighing 45 ± 3.8 kg (mean ±SD). All procedures described were performed on the right cervical VN in acute terminal studies under isoflurane anesthesia and lasted 10-12 hours. We recorded heart rate (HR), blood pressure (BP), saturation of peripheral oxygen (SpO_2_), blood glucose levels (BGL), end-tidal carbon dioxide (CO_2_) via capnography, electroneurogram (ENG) of the VN via longitudinal interfascicular electrodes (LIFE), and electromyogram (EMG) of deep neck muscles. These recordings were obtained both as data for analyses and to ensure the welfare of the animal.

### Materials

#### Recording Devices: Longitudinal Intrafascicular Electrodes (LIFE)

Electroneurogram (ENG) recordings were conducted using LIFE devices similar to previous work (Malagodi et al., 1989; Nicolai et al., 2020). The recording electrodes were cleaned and reused between experiments, with repairs made, or electrodes remanufactured, as needed.

A DIN 42802 touch-proof connector (Digitimer, Welwyn Garden City, United Kingdom) was soldered to a stainless-steel wire (SS) (ID: 0.001 in, OD: 0.0055 in, 10-15 cm length), which was soldered to a single strand platinum iridium (PtIr) wire (OD: 0.004 in, ID: 0.002 in) using special flux (Worthington Stainless Steel Flux, Mfr. Model #331929, Columbus, OH). Heat shrink tubing was applied to the connections and silicone was applied to the ends of the heat shrink tubing to protect the solder joints from adverse environmental effects and to stabilize the position of the protective tubing. A recording window was created in the PtIr wire by removing approximately 1 mm of insulation using a thermal wire stripper (Patco Services Inc, PTS-30S). This recording window was approximately 2 cm distal to the SS/PtIr solder joint. Using 7-0 silk suture, a surgeon’s knot was tied around the PtIr wire 2 mm from the window (toward the SS/PtIr solder joint). The suture was covered in Quick-Sil (WPI, Sarasota, Florida) to serve as a mechanical stop and ensure that the exposed electrode window was within the nerve.

#### Recording Devices: EMG

EMG recordings were obtained using Rhythmlink bipolar stainless-steel needle electrodes (RLSND121-2.5, Columbia, SC).

#### Recording Devices: Additional Equipment

Tucker Davis Technologies systems (Alachua, FL; W8, IZ2MH, RZ5D, RZ6, PZ5, and SIM) were used to control stimulation and to record EMG and ENG signals. Physiological recordings, including invasive arterial pressure (Millar Inc., Houston, TX, Model #SPR-350S), were captured via AD Instruments PowerLab 8/35 and visualized in real time via LabChart (ADInstruments, Sydney, Australia). Vecuronium was delivered via Medfusion 3500 Syringe Infusion Pump (Cary, NC) to achieve skeletal muscle paralysis (dose rates in Methods: Experimental Protocol).

Anesthesia was provided via Landmark Veterinary Anesthesia Machine VSA-2100 (Louisville, KY) with Midmark Matrx Model 3000 ventilator (Dayton, OH) and SurgiVet SOMNI 3 vaporizer (Plymouth, MN).

#### Stimulating Device: ImThera

The ImThera is a medical device that features six independent stimulating electrodes or “contacts,” initially designed to stimulate the hypoglossal nerve (Figure 1). To avoid ambiguity, the ImThera will be referenced as a “device” or “cuff” throughout this paper, while the six independent stimulating electrodes will be referred to as “electrodes” or “stimulating contacts.” When implanted, the device forms a cuff around the nerve, allowing its contacts to have axial and longitudinal separation along the nerve (Figure 1).

**Figure 1.**
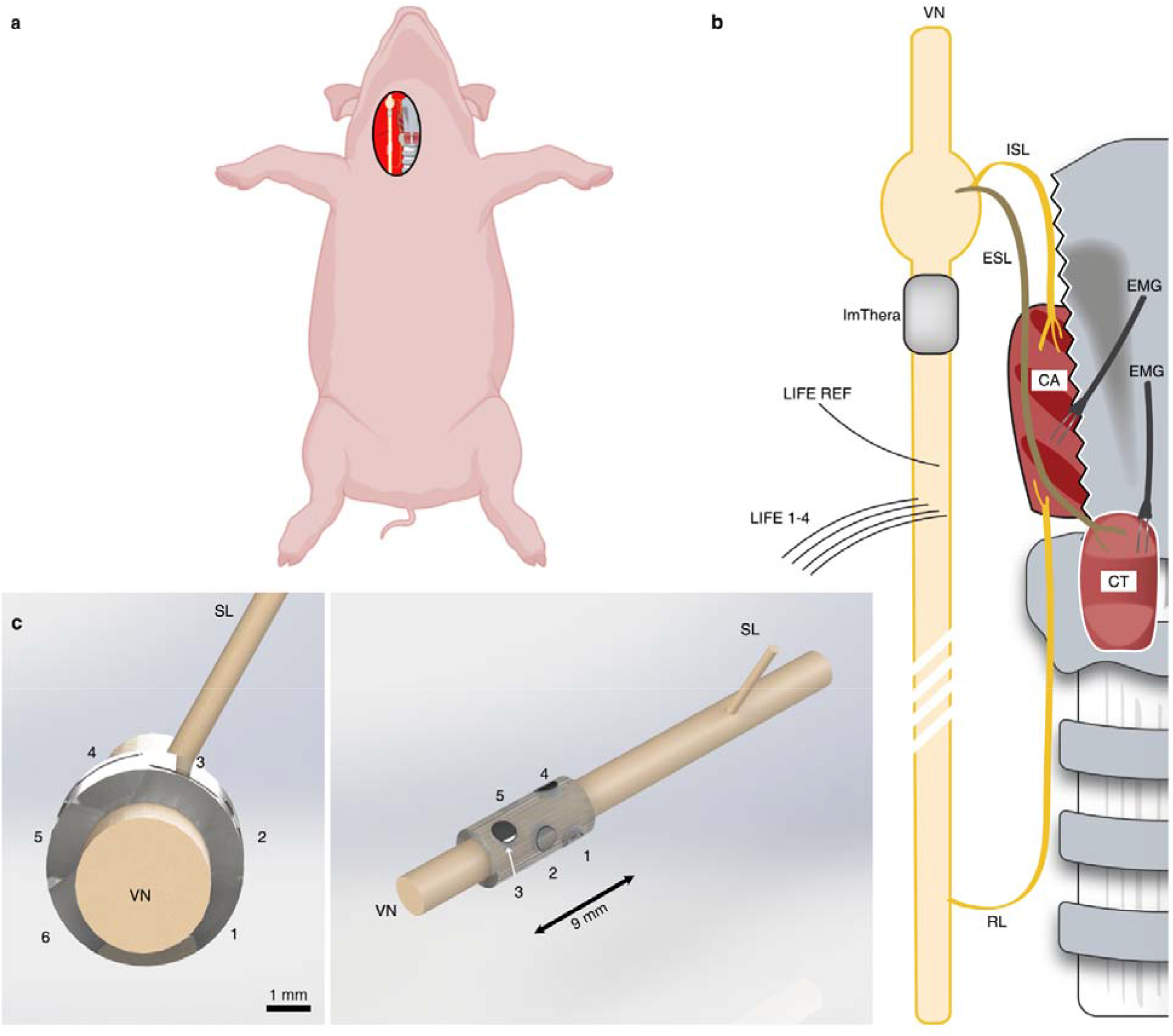
Experimental and instrumentational overview. **a,** Depiction of the surgical preparation in which the right cervical vagus nerve (VN) is exposed. **b**, Depiction of pertinent anatomy. The superior laryngeal (SL) nerve branches into the internal superior laryngeal (ISL) nerve that innervates the cricoarytenoid (CA) muscle and the external superior laryngeal (ESL) nerve that innervates the cricothyroid (CT) muscle. The recurrent laryngeal (RL) nerve also innervates the CA. “Direct” activation of the neck muscles occurs via activation of motor nerve fibers from the stimulating electrode, branching at the RL and innervating the CA. “Indirect” activation of the neck muscles results from current escaping the stimulating cuff, traveling through fluid and surrounding tissue and activating motor nerve fibers in the SL, which causes contraction of the CA and CT. This figure is not drawn to scale, but the representation of a longer RL than SL is accurate and an essential factor affecting EMG latencies. **c(left),** Transverse plane depiction of the six-contact ImThera device on the VN. The caudal direction points out of the page, towards the reader. **c (right),** Additional perspective of the ImThera cuff, showing the axial and longitudinal orientation of stimulating contacts.

### Methods

#### Validating Electrochemical Properties of Recording and Stimulating Devices

As outlined by Wilks *et al.* (Wilks et al., 2017), voltage transient (VT) analysis, electrochemical impedance spectroscopy (EIS), and cyclic voltammetry (CV) were conducted to document the properties of the electrodes (see Supplementary Material: Electrochemical Testing and Validation). Briefly, EIS was performed on all devices prior to each surgery for quality control. LIFE devices with an impedance below 10 kΩ at 1 kHz were deemed suitable and used in the experiment. All ImThera EIS measurements were stereotypical of platinum-iridium electrodes (see Supplementary Material: Figure 1). Additionally, VT and CV analyses were conducted upon receipt of the ImThera to establish safe stimulation limits (oxidation/reduction and charge injection limits) (Supplementary Material: Electrochemical Testing and Validation).

#### Surgical Methods: General Overview

All animals (n = 6) were sedated using an intramuscular injection of Telazol (6 mg/kg) and xylazine (2 mg/kg). The animals were then intubated and anesthetized using 1-2% isoflurane. Depth of anesthesia was routinely assessed (via palpebral reflex or nasal septum pinch) and drug dosages were appropriately adjusted to ensure proper sedation levels. Respiration was controlled via a ventilator set to 13-18 breaths per minute to keep end-tidal CO_2_ levels within 34-45 mmHg. Normal saline was administered continuously via IV with analgesia (Fentanyl, initial 5 μg/kg bolus, 3-5 μg/kg/hour maintenance). Heart rate was measured via EKG. Pulse rate and SpO_2_ were measured via an infrared (IR) probe placed on either the tongue or lip. BP was measured via an invasive probe (Millar) placed in either the left or right femoral artery. The final three animals additionally had their BGL values monitored every 30-60 minutes. Animals were euthanized using IV KCl (4 mEq/kg).

An incision was made via electrocautery lateral (right) of midline, between the mandible and sternal notch, to access the right VN. In two animals, the omohyoid muscle had to be transected to reach the carotid sheath. Blunt dissection was used to isolate the VN from the carotid artery within the carotid sheath. The VN was exposed from the inferior (nodose) ganglion to approximately 10 cm caudal of this point. The length of the surgical window was intended to maximize the distance between stimulating and recording electrodes to allow separation between the stimulus artifact and the fast-traveling electrically evoked compound action potentials (eCAPs). Care was taken to avoid applying mechanical forces on the VN while other structures were isolated using surgical vessel loops. The surgical window was kept open throughout the procedure. Pooled liquid in the surgical window was removed periodically with manual wicking. Saline-soaked gauze was placed around exposed portions of the VN, and saline spray was applied to the surgical window as needed to keep all structures tissues hydrated.

#### Surgical Methods: ImThera Implantation

The proximal edge of the ImThera device was placed on the right VN caudal to the nodose ganglion at a distance of 1.1 ± 0.80 mm. A 7-0 silk suture was placed around the device to keep it stationary. Effort was made to minimize mechanical forces on the VN from the device.

#### Surgical Methods: LIFE Implantation

Five LIFE devices were placed within the VN caudal to the ImThera cuff to record ENG, caudal to the ImThera cuff. These devices were implanted longitudinally (caudal to cranial) such that the entirety of the recording window was parallel to the course of the nerve. The most cranial LIFE served as a reference electrode for the remaining four LIFE devices. The recording electrodes were staggered in the cranial/caudal direction to discern spatiotemporal aspects of the ENG signals. Cranial/caudal staggering ensured signals were confirmed as neural in origin if temporal differences were observed among recording electrodes. Medial/lateral staggering enabled sampling and later analysis of ENG signals from various locations within the nerve. The distance between the stimulating electrode and the recording electrodes was maximized within the surgical window to reduce the contamination of the neural signals with stimulation artifact. This resulted in an average distance of 7.93 ± 1.75 cm from the caudal edge of the cuff to the most cranial LIFE and 0.79 ± 0.38 cm from the most cranial to the most caudal LIFE.

#### Surgical Methods: EMG Implantation

EMG needle electrode pairs were placed in the cricothyroid, cricoarytenoid, and pharyngeal constrictor muscles.

#### Experimental Protocol

We conducted stimulation trials in the “intact” condition, followed by multiple control conditions. In all cases we stimulated to approximate clinical parameters, using a symmetric biphasic rectangular pulse with 0 ms interphase delay, 0.2 ms per phase, cathodic phase first, and 0.4 ms total duration, mimicking common clinical parameters (Supplementary Material: Figure 2) (Koo et al., 2001). All stimulating pulses were delivered in a monopolar configuration to avoid potential interactions between simultaneously active cathode/anode electrode pairs along the nerve. We placed an 18-gauge, stainless steel needle (152 mm^2^ surface area) within the thoracic wall to serve as the counter electrode. The stimulus was delivered through one of six contacts of the ImThera cuff at 25 Hz with random amplitudes from 50 to 5000 μA. We paused for 60 s between each amplitude to allow for physiological states to return to baseline. The range of randomized stimulation amplitudes was adjusted on a case-by-case basis based on physiological responses (e.g., in some animals, severe bradycardia was evoked during stimulation at high amplitudes, see Results). Stimulation was delivered sequentially through electrode contacts 1 through 6. For each contact and stimulation amplitude, a total of 750 pulses (i.e., 30 s of stimulation) per amplitude per stimulation contact were delivered in the intact condition. Subsequently, animals were stimulated under the following conditions to confirm the source of recorded signals: 1) neuromuscular junction (NMJ) block (vecuronium, IV, 0.15 mg/kg bolus, 0.15 mg/kg/hour maintenance rate), 2) transection of the superior laryngeal (SL) branch, 3) transection of the recurrent laryngeal (RL) branch, and 4) transection of the vagus trunk cranial and caudal to the stimulating cuff (double vagotomy). Multiple studies demonstrated that EMG signals can appear as artifacts in neural recordings (Nicolai et al., 2020; Yoo et al., 2013); therefore, we conducted both NMJ block and transections to assess the origin of recorded signals (Supplementary Material: Off-Target Activation). After stimulating in the NMJ condition, we waited for the paralytic agent to wear off before proceeding with the remaining conditions. In all these additional conditions (NMJ to double vagotomy), we delivered 25 pulses per amplitude (i.e., 1 s of stimulation) due to limitations in our protocol-allotted surgical time.

**Figure 2.**
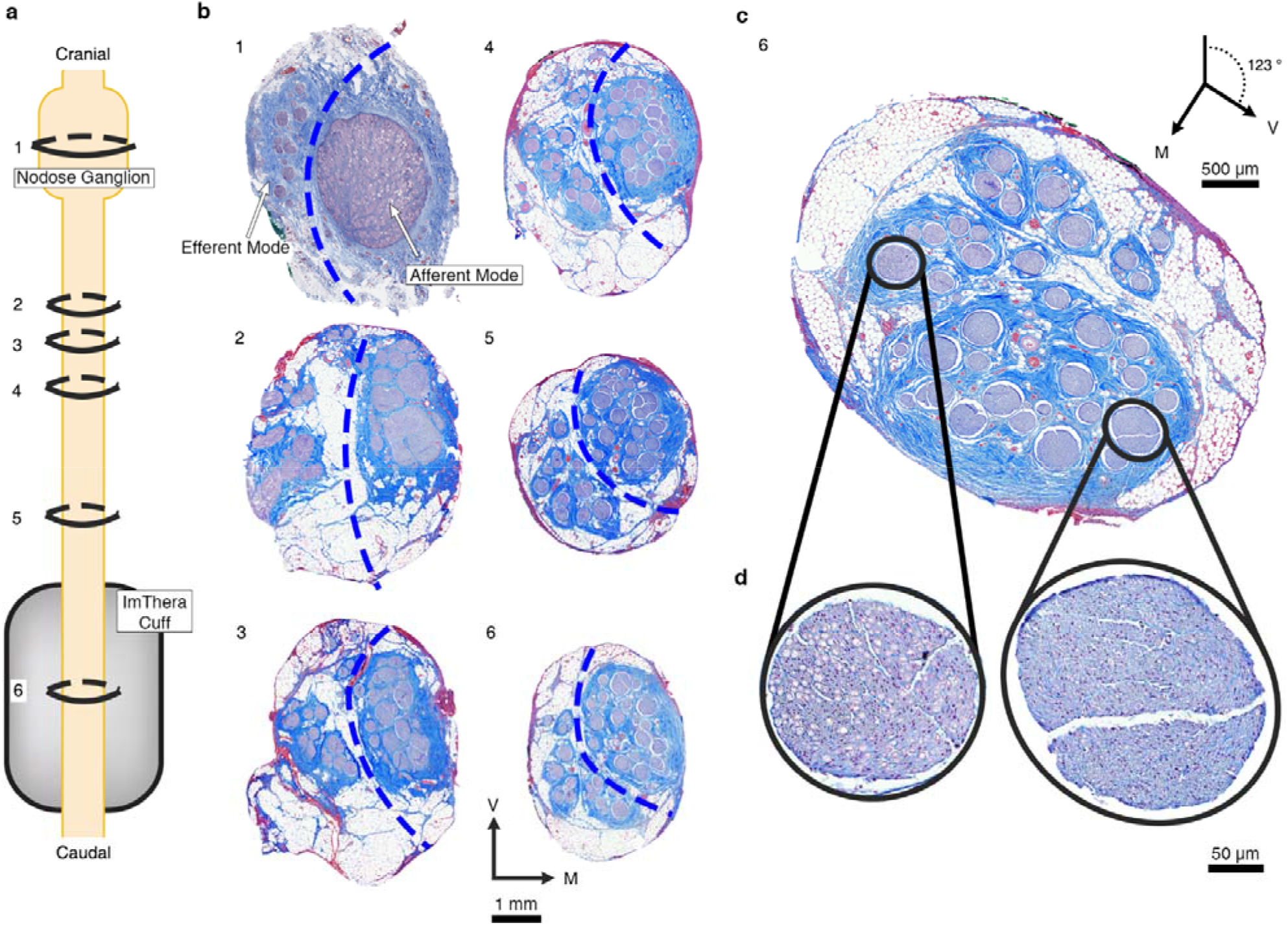
Histology from the nodose ganglion to the level of the stimulating cuff showing bimodal distribution of efferent and afferent fascicles in the vagus nerve of a pig. **a,** Depiction of the VN from nodose ganglion (known point of organization of afferent fascicles) to the level of the cuff. Rings indicate the proportional locations of histological cross sections 1-6 in panel b (spanning 1.04 cm total, from slice #1 to slice #6) for Animal #4. **b**, Six histological cross sections of the VN for Animal #4. The serial sections track the locations of the fascicles arising from the pseudounipolar cell bodies of the visceral afferent fibers in the nodose. The dashed blue lines were superimposed to segregate the efferent and afferent modes based on the separation of the soma and the efferent fibers of passage in slice #1; we then followed these afferent and efferent groupings caudally. The most caudal section (6) is repeated in **panel c** and rotated 123° clockwise to place the efferent mode in the top half of the section. **d**, The zoomed views shows the large, myelinated fibers in an example efferent fascicle (left) and the small, unmyelinated fibers in an example afferent fascicle (right). V: ventral; M: medial.

#### Histological Analysis

Histology of the nerve was conducted to relate vagotopy with stimulation-evoked physiological responses. To ensure that no movement had occurred during the procedure, the placement of the cuff on the nerve (longitudinal location and relative rotation) was recorded and measurements to known landmarks (e.g., nodose ganglion) were made before and after the procedure. Following all functional studies and euthanasia, animals underwent microdissection and histological procedures as previously described (Settell et al., 2020). Briefly, the VN was exposed from the nodose ganglion to the recurrent laryngeal bifurcation at the level of the subclavian artery. To match histology to relative *in vivo* locations, tissue dye was used to mark the ventral and lateral edges of the nerve (Bradley Products, Inc., Davidson Marking System, Bloomington, MN), and sections were removed for fixation and processing. All sections were placed in 10% neutral buffered formalin at 4 °C for 24-48 hours. Samples then were embedded in paraffin wax and 5 μm thick slices, approximately every 40 μm, were collected and mounted on charged slides (Sakura Tissue-Tek VIP). Slices were stained with Gomori’s trichrome and were imaged with a Motic Slide Scanner (Motic North America, Richmond, British Columbia). Measurements acquired during the surgery were used to superimpose the *in vivo* location and orientation of the cuff onto the histology.

#### Data Analysis

##### Preprocessing ENG and EMG Signals

The ENG and EMG data were digitally filtered using in-house developed software (Ludwig Lab, 2021/2021) to increase the signal-to-noise ratio and to remove long-term baseline drift without introducing filter ringing from a large stimulation artifact, which could otherwise be misconstrued as neural signals (Lindeberg, 1990; Widmann & Schröger, 2012). Specifically, we subtracted the median filtered signal (kernel size: 201) from the original signal (Jarske & Vainio, 1993). Next, we applied a Gaussian filter (standard deviation for Gaussian kernel: 0.87) and a finite impulse response filter to reject common-mode noise and its harmonics (60, 120, and 180 Hz) with 1.0 Hz bandwidth. An additional median filter (kernel size: 11) was applied to EMG signals to eliminate intermittent spikes present only in EMG recordings.

##### Evoked Compound Action Potential Magnitude Calculation

After filtering, neural signals were averaged across pulse trains and segmented into five time-restricted windows based on known conduction velocities per fiber type (i.e., Aα, Aβ, A⍰, Aδ, and B) (Supplementary Material: Table 1). The onset of stimulation was set as the time zero starting point. The boundaries of the time-restricted windows were calculated by diving the distance from the stimulating device to the recording electrode by the conduction velocity of each fiber type. For neural signals uncontaminated by stimulation artifact, the root mean square of the measured voltage (V_RMS_) within each time-restricted window was calculated to determine the magnitude of the eCAP generated by that nerve fiber type.

If neural signals contained noticeable eCAPs but were still contaminated by stimulation artifact despite the preprocessing steps, the signals were processed additionally to isolate the eCAP (Supplementary Material: Figure 3). First, to find the onset and offset of the eCAP, the contaminated signals were filtered using a Savitzky-Golay filter to smooth the signal and eliminate false minima (Savitzky & Golay, 1964). Next, a peak-detection algorithm was applied to find the local maximum within the time window for each fiber type. The time points of the closest minima occurring before and after the maximum were classified as onset and offset of the eCAP. The later minimum (offset) could exceed the bounds of the time-restricted window by 25% to allow for the entirety of the signal to be captured. Once the onset and offset of the eCAP were found, these points were superimposed on the filtered signal (without Savitzky-Golay filter). Similar to other studies examining the amplitude of the evoked response in EMG recordings by calculating the integral of the measured signal (Haughton et al., 1994; Kent-Braun, 1999; Zory et al., 2005), we calculated the definite integral of the eCAP. To account for the stimulation artifact, a simplified model of the stimulation artifact was created by forming a line between the onset and offset of the eCAP. The definite integral of this simplified model was subtracted from the definite integral of the signal to give an area measurement of the eCAP (Supplementary Material: Figure 3).

**Figure 3.**
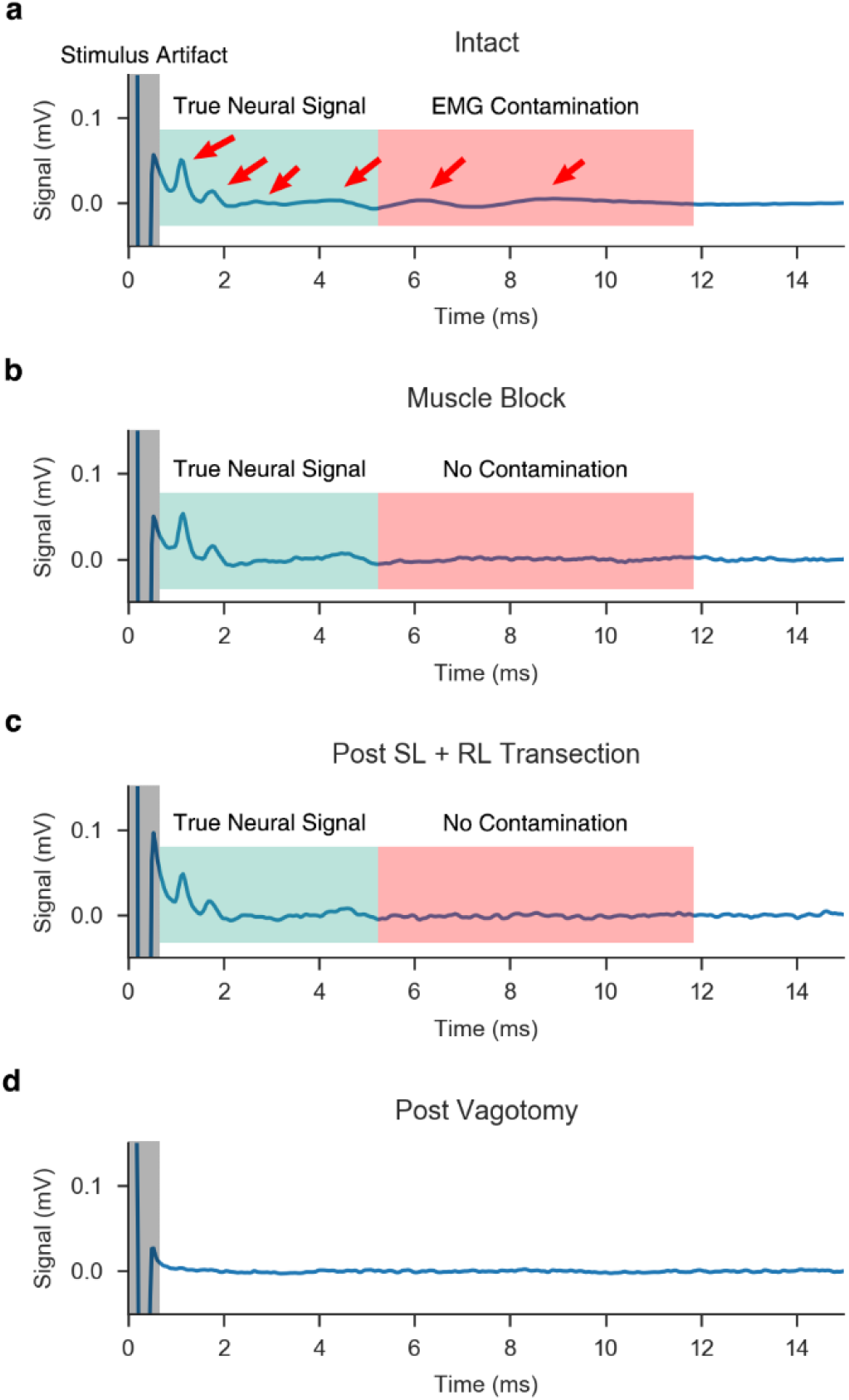
Representative ENG signals measured by intrafascicular electrodes. **a,** Signal measured via LIFE in the VN during the “intact” condition. Candidate eCAPs are denoted by red arrows. Candidate eCAPs in the green shaded region correspond to the latencies of Aα and B fibers, while candidate eCAPs in the red shaded region correspond to the latencies of Aγ to B fibers. In this condition, it is difficult to determine which signals are true eCAPs and which signals are EMG contamination. Also, note the stimulus artifact at the onset of the stimulation (grey shaded region). **b**, Upon applying a paralytic agent, the EMG is eliminated and only the neural signal remains in the recorded trace. **c,** Without paralytic, RL and SL branches are transected, eliminating direct and indirect pathways to nearby muscles within the neck, which provides an additional confirmation of the lack of EMG contamination in the ENG signal. **d**, Verification of neural signal sources via double transection of the vagus nerve, rostral and caudal to the stimulation cuff, after which no signal remains. Note, the “intact” condition (panel a) is the result of averaging the recordings from 750 pulses, while the remaining conditions (panels b-d) are averaged across 25 pulses, resulting in slightly more noise in the traces.

##### Dose-Response Curve Calculation

Dose-response curves (DRCs) were generated for neural, muscle, and physiological responses to randomized stimulation amplitudes. The “intact” condition was used to analyze: 1) nerve fiber responses measured via ENG, 2) muscle response measured via EMG, and 3) changes in HR and BP from baseline measured via EKG and invasive arterial catheter, respectively.

As described above, the magnitude of the eCAPs was calculated either as V_RMS_ or as area-under-the-curve following the peak-detection algorithm. The amplitude of the EMG response was calculated via V_RMS_; as evoked EMG responses occurred outside of the window contaminated by the stimulus artifact (~5 ms), no stimulus subtraction methods were required to measure the magnitude of the evoked signal. We calculated 95% confidence intervals (CI) for EMG and ENG responses by bootstrapping the sampled signal and calculating the amplitude of the measured eCAP based on the bootstrapped sample estimates (see Supplementary Material: Interpolation and Statistics). Changes in HR and BP were defined as the maximum change during a stimulation pulse train from the baseline calculated over the interval immediately prior to stimulation (i.e., t-3 to t-1 seconds before stimulation).

##### Response Interpolation and Statistical Analyses

We examined trends in evoked measurements based on the relative location of the stimulating electrode to the efferent and afferent clusters within the VN. For each animal, the cross-sectional histology of the VN (at the location of the center of the stimulation cuff) was annotated with the locations of the six stimulating contacts (Supplementary Material: Figure 4a). To ensure that the impact of electrode location with respect to the efferent mode was consistent across the cohort analyses, the histology/contact figure was rotated such that the efferent-mode fibers were in the top half of each image (12 o’clock position) and the afferent-mode fibers were in the bottom half (6 o’clock position) (Supplementary Material: Figure 4b).

**Figure 4.**
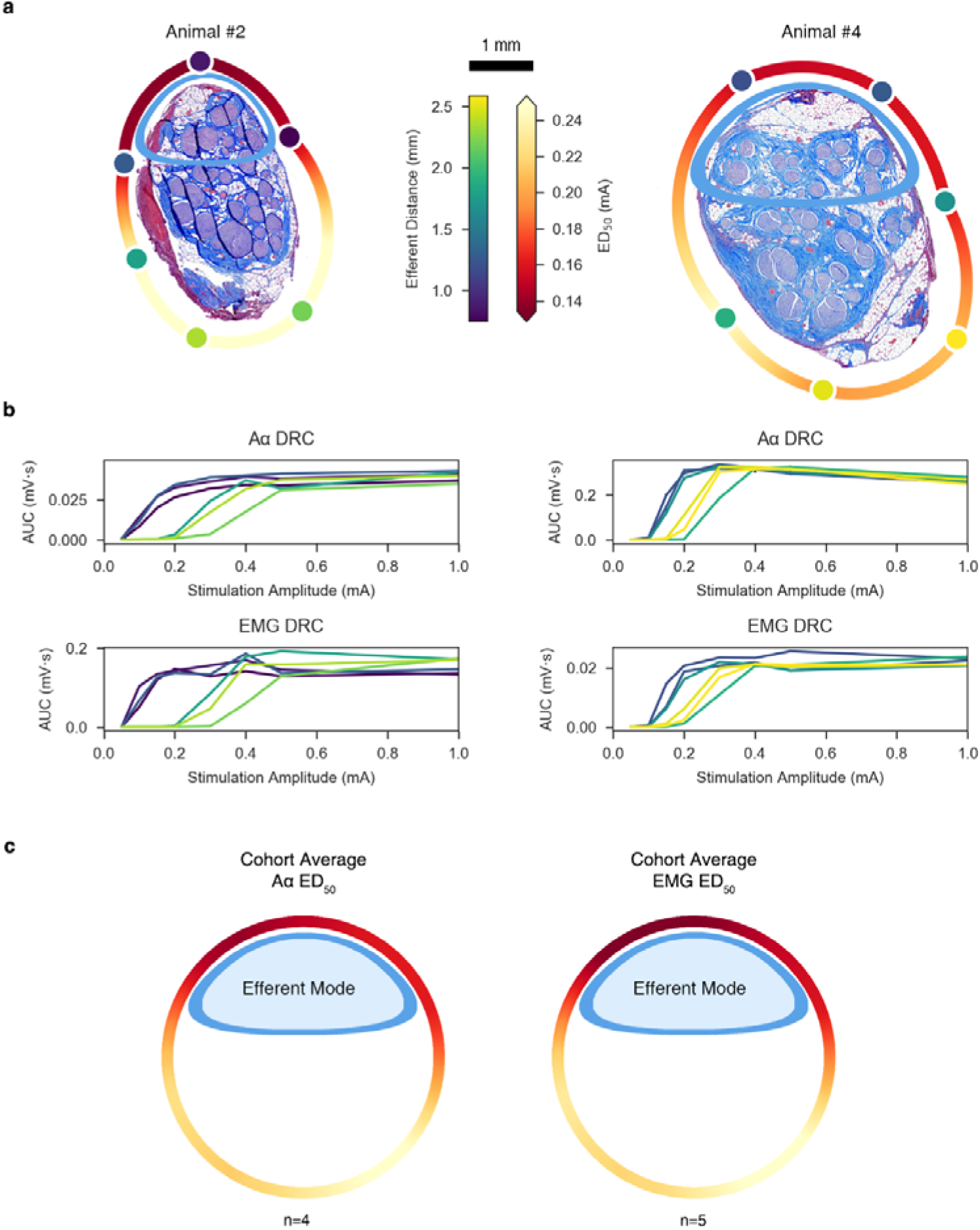
ENG and EMG dose-response curves reflect contact position-dependent responses. **a,** Histology for Animal #2 (left) and Animal #4 (right) at the level of the stimulating cuff, rotated such that the efferent cluster is oriented toward the top of the page. Stimulating contacts are superimposed to replicate the *in situ* condition and are characterized in color by their distance to the centroid of the efferent cluster (“Efferent Distance” colorbar). A heatmap was interpolated around the perimeter of the histology to indicate the current amplitude required to elicit half-maximal Aα responses (“ED_50_” colorbar). **b,** ENG and EMG dose-response curves for Animal #2 (left) and Animal #4 (right); 95% CIs are plotted but not visible due to its extremely narrow range^2^. The color of each trace corresponds to the colors of the contacts in panel a. **c,** Extrapolated ED_50_ heatmap for both neural (left; n = 4) and muscle (right; n = 5) responses, averaged across the cohort and superimposed on a generalized circular illustration of the nerve cross section. The smaller ellipsoid (blue) represents the efferent mode. Same colorbar for ED_50_ as in panel a.

In each animal, for each of the six stimulating contacts, we calculated the maximum change in HR and the stimulation amplitudes that elicited 50% of the maximum response (ED_50_) for Aα and EMG. To determine ED_50_ values for Aα and EMG measurements, we fit the neural and EMG DRCs to the sigmoidal Hill equation (A.V. Hill, 1910) using GraphPad (Prism, San Diego, CA). For each metric (Aα ED_50_, EMG ED_50_, maximal HR change) in each animal, we linearly interpolated 500 values between the neighboring contacts along the nerve circumference, which resulted in 3000 values (500 data points x 6 neighboring contact pairs) per metric, per animal. These 3000 values were plotted around the perimeter of the rotated micrograph (efferent-mode facing the 12 o’clock position) using Python.

To build an aggregate heatmap and calculate statistics across the cohort, a generalized VN cross-section was created utilizing the same efferent/afferent oriented gestalt, and the interpolated values were averaged, based on their relative location, across the cohort. For the cohort maximum change in HR, Aα ED_50_, and EMG ED_50_, we averaged the interpolated arrays across the animals for which we were able to obtain recordings with sufficient resolution (n = 6, n = 4, n = 5, respectively).

Additionally, we conducted a paired t-test comparing the Aα ED_50_ values, across the cohort, for the three contacts closest to the efferent cluster versus the ED_50_ values for the three contacts furthest from the efferent cluster.

#### Computational Modeling

We used the ASCENT (“Automated Simulations to Characterize Electrical Nerve Thresholds”) pipeline v1.1.0 (Musselman et al., 2021) to model VNS for all six pigs with the ImThera cuff using individual-specific nerve morphology from histology (Supplementary Material: Figures 5 and 6, Table 2). For each pig, we segmented the epineurium and fascicles in the histological cross section at the level of the ImThera cuff using (NIS-Elements Ar software, v5.02.01, Build 1270, Nikon Instruments Inc.) as described previously (Pelot et al., 2020). We modeled the three-dimensional geometry of the ImThera cuff around the modeled nerve, a 100 μm layer of saline around the cuff, and the surrounding tissue as muscle (25 mm length, 10 mm diameter). ASCENT built, meshed, and solved the finite element models using COMSOL Multiphysics v5.4 (Burlington, MA). For each of the solved COMSOL models—one for each of the six contacts for each of the six animals (total = 6×6 = 36 models)—the ASCENT pipeline sampled the resulting electric potentials along the length of the nerve at the centroid of each fascicle, i.e., along the paths of axons. Using NEURON v7.6 (Hines & Carnevale, 1997), ASCENT applied the potentials to biophysically-realistic models of mammalian myelinated nerve fibers, corresponding to the large-diameter (13 μm MRG) myelinated Aα and medium-diameter (3 μm MRG) myelinated B fibers (McIntyre et al., 2004; *Modeling the Excitability of Mammalian Nerve Fibers: Influence of Afterpotentials on the Recovery Cycle / Journal of Neurophysiology*, n.d.; Musselman et al., 2021). We used a binary search algorithm (1% resolution) to identify the activation threshold for each fiber in response to a charge-balanced, biphasic pulse (200 μs/phase with no interphase delay, as used in vivo). Data plotting and analyses were done in Python v3.7 (Van Rossum & Drake, 2009). Further details are provided in Supplementary Materials: Computational Modeling.

**Figure 5.**
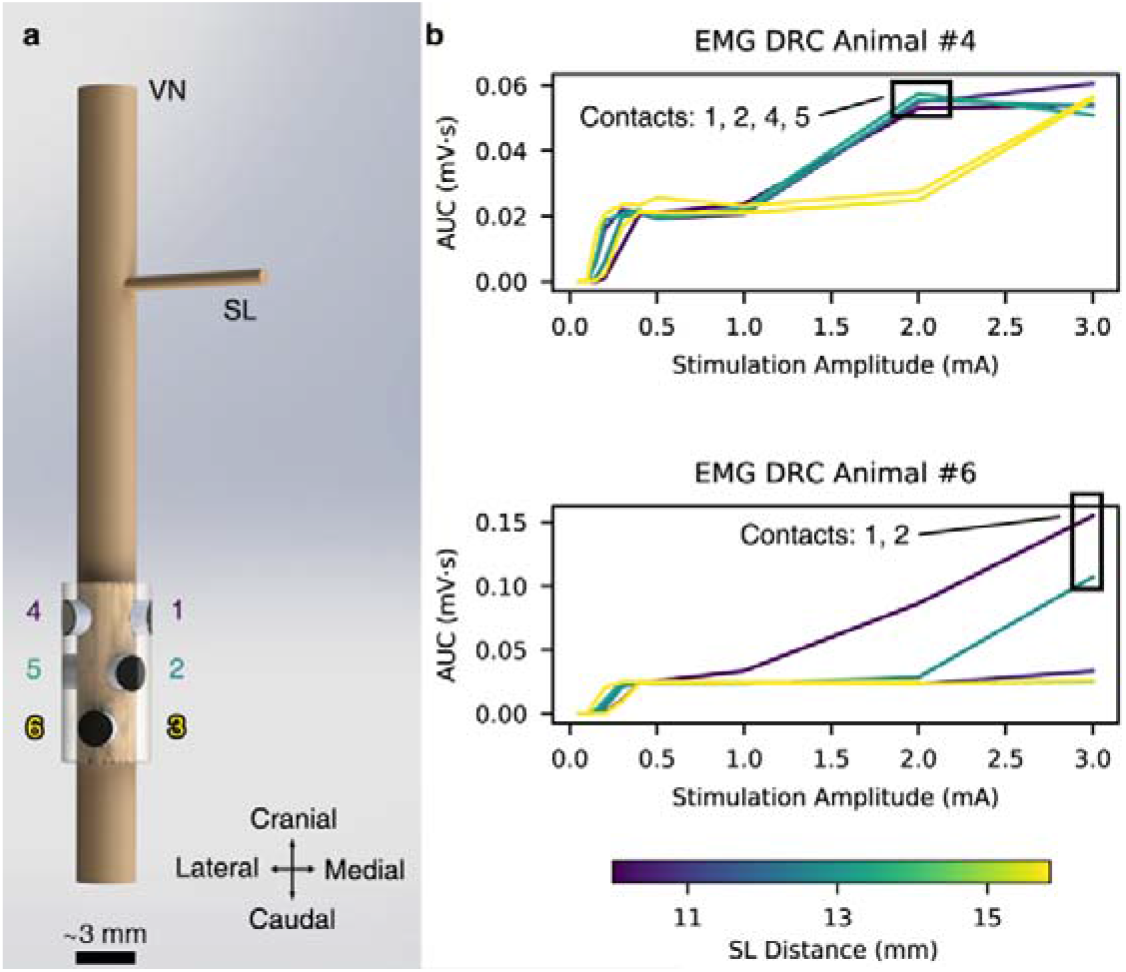
Off-target activation of the SL. **a,** 3D rendering of the ImThera cuff placed on the VN showing the locations of the stimulating contacts relative to the SL branch. **b,** Dose-response curves (DRCs) in the animals with additional EMG recruitment at stimulation amplitudes >1 mA. Contacts closest to the SL branch preferentially activate the SL fibers at lower stimulation amplitudes than those that are further away.

**Figure 6.**
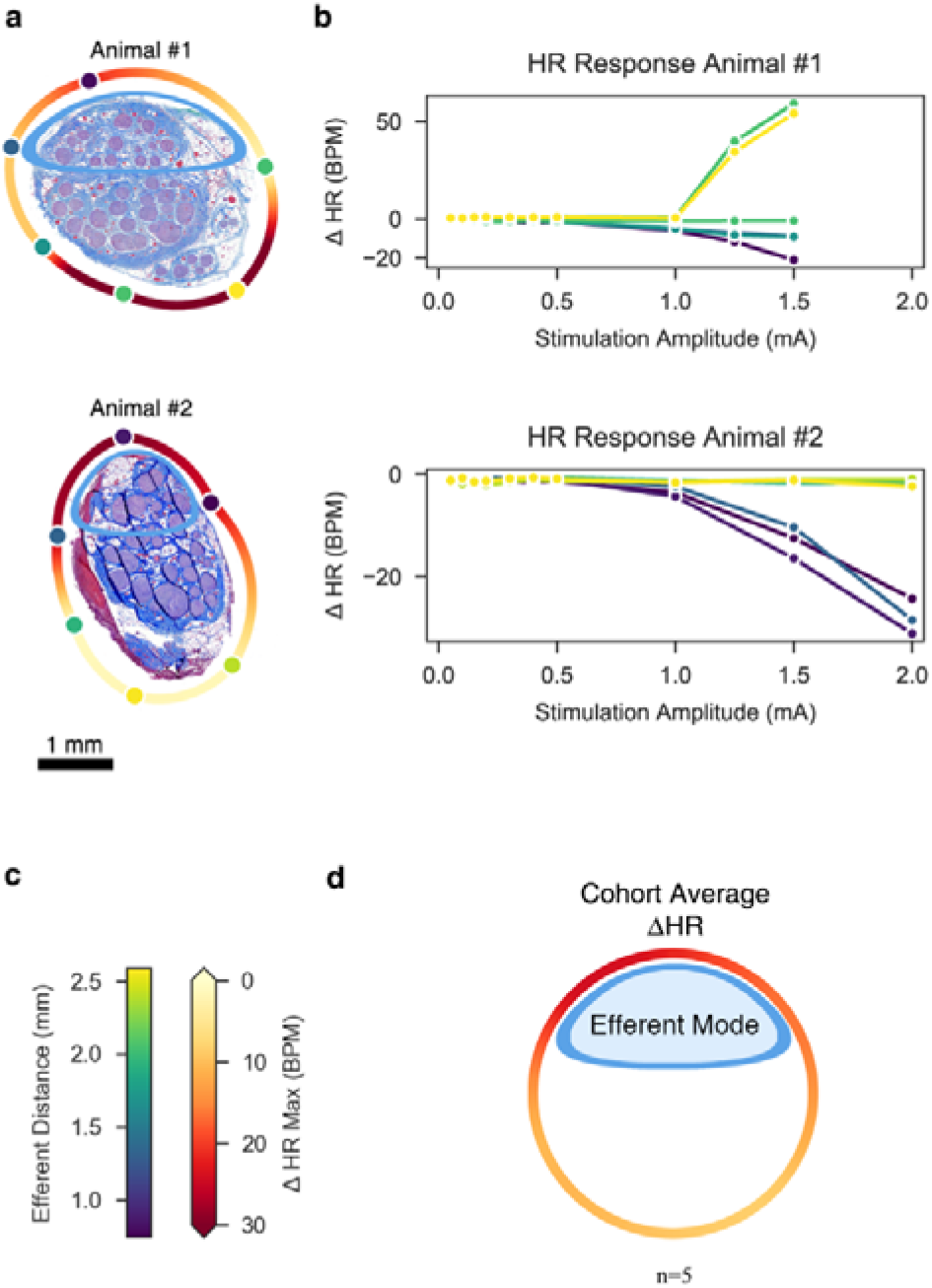
Heart rate dose-response curves reflect electrode contact position-dependent recruitment. **a,** Histology for Animal #1 (top) and Animal #2 (bottom) at the level of the stimulating cuff, rotated such that the efferent cluster is oriented toward the top of the page. Stimulating contacts are superimposed to replicate the *in situ* condition and their colors indicate their distances to the centroid of the efferent cluster. A heatmap was extrapolated around the perimeter of the nerve to indicate the maximal change in HR, based on location (see colorbar in panel c). **b,** Heart rate (HR) dose-response curves. Colors match the stimulating contacts in panel a. Note the marked tachycardia (increased HR) in Animal #1, particularly for contacts closest to the additional small cluster of fascicles at the bottom right of the histology. Conversely, both animals show bradycardia (decreased HR) for contacts near the efferent cluster at the top of each micrograph. **c,** Colorbar legends for the distance of each contact to the efferent cluster and for the maximum change in HR. Note that these legends are not related to each other (e.g., 2.5 mm distance does not correspond to 0 bpm change in HR). **d**, Extrapolated heatmap of the change in heart rate, averaged across the cohort (n = 5) and superimposed on a generalized circular illustration of the nerve cross section.

We used a mixed model ANOVA for each fiber type to assess factors that influenced the models’ relative error to in vivo thresholds (fixed effects: full factorial of vagotopy (included in model or not), response level (Aα fibers only), distance from the efferent mode; random effect: animal). For significant terms, we performed paired student’s t-tests on the models’ errors with and without the fixed effect.

## Results

The goal of this study was to understand how the underlying anatomy of the pig cervical VN relates to functional response to VNS evoked using an FDA IDE-approved six-contact cuff electrode, with a focus on developing a conceptual framework for differentiating putative therapeutic effect (i.e., heart rate reduction) from side effect (concomitant laryngeal muscle activation). Data from the recent NECTAR trial of VNS to treat heart failure suggested that the intended changes in sympathetic/parasympathetic tone were limited by unwanted neck muscle activation (De Ferrari et al., 2017); therefore, we used the stimulation-evoked changes in heart rate as a primary measure of therapeutic engagement. In the following sections, each animal’s histology is presented first to provide context for evaluating the observed functional results. Next, the physiological results are presented in context of the underlying fascicular organization and the associated positions of the active electrode contacts. Finally, subject-specific computational modeling is used to develop a conceptual framework to understand how the underlying fiber type-specific fascicular organization relates to the observed functional results.

### Histology to Trace Efferent and Afferent Fascicles from Nodose to Surgical Window

The vagus nerve has distinct functional organization at specific points along its path connecting the brainstem to the visceral organs (Câmara & Griessenauer, 2015; Ellis, 1964). Somatic motor efferents responsible for deep neck muscle activation originate within the nucleus ambiguus in the medulla oblongata and more distally become the pharyngeal, superior laryngeal (SL), and recurrent laryngeal (RL) branches, which innervate the pharyngeal, cricothyroid (CT), and cricoarytenoid (CA) muscles of the deep neck (Figure 1). Parasympathetic efferents originate in the dorsal motor nucleus of the vagus in the medulla oblongata and distally become vagal branches to visceral organs. Visceral afferents course within these same branches back to the cervical vagus; these afferents are pseudounipolar cells with cell bodies in the nodose ganglion and central axons terminating in the solitary nucleus (NTS).

Across the cohort, histology of the VN at the level of the cuff showed a functionally clustered organization of fascicles (Supplementary Material: Figure 4). Cranial to the cuff, pseudounipolar cell bodies aggregate in the nodose ganglion (Figure 2b, slice 1) (Settell et al., 2020). These afferent neurons of the nodose remained clustered in a distinct grouping of fascicles as they descended caudally to the level of the surgical window (Figure 2b, slices 2-6, “Afferent Mode”). We traced this organization by taking histological slices every ~40 μm. The remaining fascicles (Figure 2b, “Efferent Mode”) contained both motor and parasympathetic efferents^1^ (Câmara & Griessenauer, 2015; Ellis, 1964; Rea, 2014; Settell et al., 2020). Thus, the cross section of the nerve at the level of the stimulation cuff contained two distinct functional groupings of fascicles: an efferent grouping and an afferent grouping (Figure 2b, slice 6). Further, the efferent grouping of fascicles is expected to preferentially contain the largest diameter myelinated motor nerve fibers while the afferent fascicle cluster is expected to contain smaller diameter myelinated and unmyelinated sensory afferents from the visceral organs; indeed, we observed this distinction in fiber diameters between fascicles in both modes to confirm the presence or relative absence of the largest diameter myelinated fibers (Figure 2d).

### Sources of Electroneurography and Electromyography Signals

We conducted multiple controls to confirm the sources of the various components of the EMG and ENG signals. We first considered the relevant gross anatomy, including the neural pathways activated by VNS that evoke EMG responses: the RL and SL branches of the vagus (Figure 1). Prior studies demonstrated that electrical stimulation of the cervical vagus with an epineural cuff can activate motor nerve fibers within the cervical vagus that become the RL (DuBois & Foley, 1936; Foley & DuBois, 1937; Nicolai et al., 2020). In addition, motor nerve fibers of the SL which are located near the cuff can be activated through current leakage (Nicolai et al., 2020). Across the cohort, the motor nerve fibers in the RL traverse a long path (22.89 ± 1.03 cm) from the implanted cuff electrode to their innervation of the CA. Conversely, the motor nerve fibers of the SL travel a much shorter path: the external superior laryngeal (ESL) branch travels 4.7 ± 0.6 cm from the nodose to the cricoid cartilage, under which it inserts in the CT muscle, and the internal superior laryngeal (ISL) branch travels 3.4 ±0.4 cm to its insertion in the CA muscle. Activation of the long RL pathway causes muscle contractions, and the resulting EMG appears as artifact in electroneurography (ENG) recordings that could easily be conflated with activation of fibers with lower conduction velocities, i.e., long latency components in the evoked compound action potential (eCAP) (Figure 3 and Supplementary Material: Figure 20) (Nicolai et al., 2020). Activation of the SL pathway generates a shorter latency EMG response that could also appear as artifact in the ENG recordings (Supplementary Material: Figure 20, 21).

We confirmed the sources of the eCAP versus EMG artifact in the ENG signal through use of a paralytic agent vecuronium and multiple nerve transections. As shown in Figure 3a, there were multiple peaks in the eCAP signal in the intact condition in the latency ranges for the Aα and B fibers. Transection of the RL and SL branches eliminated the two slowest peaks (Figure 3a, two right most red arrows), thus indicating their source was EMG contamination. However, transections of the RL and SL alone do not account for the possibility of an unanticipated motor nerve fiber pathway being activated and evoking EMG that appears as artifact in the ENG. Therefore, eCAPs were also obtained following intravenous administration of the muscle blocker vecuronium to eliminate all other confounding muscle signals. eCAP components after RL and SL transection or under a general muscle block remained unchanged (Figure 3b vs. c), indicating the RL and SL transections successfully eliminated all EMG contamination from the ENG recordings. Finally, we confirmed that remaining eCAP components were truly neural signals by transecting the vagus on both sides of the stimulating cuff, which eliminated all signal (Figure 3d).

In addition to controls with a paralytic agent and with transections, we also observed correlations between ENG activity and physiological responses by comparing the dose-response curves for Aα fibers with EMG across contact locations (Figure 4). Since Aα fibers have a short latency eCAP response, Aα eCAP components could be reliably discriminated from the stimulation artifact in only four of the six animals. Activation of Aα motor nerve fibers within the cervical vagus directly cause contractions of the CA muscle via the RL (Figure 1); therefore, it is reasonable to anticipate strong correlation between Aα and EMG dose-response curves. As expected, there was strong agreement between Aα ENG and the EMG dose-response curves with respect to the onset threshold (i.e., stimulation amplitude producing the initial rise in signal), saturation threshold (i.e., minimum stimulation amplitude producing maximum signal), and differences in the responses across the six contacts (Figure 4, Supplementary Material: Figure 23).

### Location-Specific Differences for Evoking Unwanted Neck Muscle Activation

We quantified the effects of contact location around the nerve circumference on the recruitment of Aα fibers and the associated EMG responses, leveraging our mapping of the underlying nerve morphology. Figure 4b shows the Aα and EMG dose-response curves for two representative animals as a function of contact location; the data across the contacts are color-coded by the distance of the contact from the centroid of the grouping of fascicles containing somatic motor and parasympathetic efferent fibers. The dose-response curves have lower thresholds for the three contacts closest to the efferent mode and higher thresholds for the three contacts furthest from the efferent mode.

We rotated the histology for each animal such that the efferent and afferent groupings were oriented toward the top and bottom of the page, respectively, to better visualize and quantify the impact of distance from the stimulating contact to the efferent fascicles on the EMG responses across the cohort (Supplementary Material: Figure 4). Next, we calculated the current amplitude required to elicit half-maximal Aα and EMG responses (ED_50_) from the dose-response curve for each contact for each animal. We mapped these amplitudes to the location of the stimulating contacts and interpolated ED_50_ values around the nerve circumference (Figure 4a). We repeated this analysis for the data for other animals (4/6 for Aα, 5/6 for EMG); the Aα signal was contaminated by the stimulation artifact in two animals, and the EMG dose-response curve had insufficient resolution to identify clearly the points of threshold and saturation in one animal. We calculated the mean ED_50_ across animals for each location around the nerve circumference (Figure 4c). As with the individual animal data (Figure 4a), this aggregate analysis shows consistency between electrode distance from the efferent cluster of fascicles and the ED_50_ values for the Aα and EMG responses, with lower thresholds for contact locations closer to the efferent fibers. We conducted a paired t-test comparing the Aα ED_50_ values for the three contacts closest to the efferent cluster versus the ED_50_ values for the three contacts furthest. As expected, the required ED_50_ for the three contacts closest to the efferent cluster (0.147 ± 0.011 mA) were lower than the ED_50_ values for the three contacts furthest from the efferent cluster (0.237 ± 0.033 mA; *t*(5) = 3.19, p = 0.0332, paired t-test).

In two of the six animals, the EMG signal increased further in response to the highest stimulation amplitudes after initially reaching a point of apparent saturation (Figure 5, Supplementary Material: Off-Target Activation). This additional increase in EMG response reflects the appearance of a new short-latency EMG response, which was eliminated by transection of the superior laryngeal branch in proximity to the cuff (Figure 1, Supplementary Material: Figures 20, 22). These transection data confirmed that the source of this additional EMG increase was current escaping the cuff to activate the nearby and shorter superior laryngeal path innervating both the cricoarytenoid and cricothyroid muscles, similar to previous studies (Nicolai et al., 2020). Off-target activation of the SL fibers occurred at lower stimulation amplitudes for contact locations closest to the SL, which branches off at a ventral-medial angle to the main trunk (Figure 5).

### Location-Specific Differences for Evoking Changes in Heart Rate

As described above, we observed statistically significant differences in EMG responses between contact locations closest to and furthest away from the efferent mode. However, the ratio of ED_50_ from the furthest to closest contact was, on average, only approximately 60% (Figure 4; Animal #2: 0.11 to 0.41 mA, Animal #4: 0.14 to 0.29 mA). In contrast, the heart rate (HR) responses to VNS were more sensitive to stimulation location (Figure 6).

Across all animals, there was at least one stimulation contact location that evoked profound bradycardia (>10 BPM) (Supplementary Material: Figure 24). The mean stimulation amplitude required to reduce HR by more than 3 BPM from baseline across all contacts and all animals was 1.3 ±0.5 mA. The mean maximum decrease in HR (HR_max↓_) across all animals was 34 ±21 BPM (range: 11 to 71 BPM; Supplementary Material: Figure 24, Animal #3 (smallest HR_max↓_), Animal #5 (largest HR_max↓_)).

Different contacts could yield broad variations in HR responses in the same animal (Supplementary Material: Figure 24). For example, in Animal #5, contact 1 yielded a HR_max↓_ of 17 BPM at 5 mA (21% change from baseline), whereas contact 4 yielded a HR_max↓_ of 71 BPM at 4 mA (89% change from baseline). In this animal, we did not stimulate at a current above 4 mA through contact 4 given the magnitude of the bradycardic response. Conversely, other contacts did not produce such a dramatic response, even at higher stimulation amplitudes. Across all animals, the difference of HR_max↓_ between the contact locations with the smallest and largest HR_max↓_ values was 35 ± 27 BPM, indicating a profound sensitivity in HR response to electrode location.

Animal #1 uniquely had profound tachycardia or marked bradycardia depending on contact location (Figure 6b). The histology showed that the two electrodes eliciting a tachycardic response were adjacent to a tertiary small grouping of fascicles, distinct from the established afferent and efferent clusters seen in the rest of the cohort (Figure 6a). This tertiary mode was assumed to be either the aortic depressor nerve or hitchhiking fascicles from the main sympathetic trunk, both of which originate from the pseudounipolar cells of the superior cervical ganglion (SCG); the aortic depressor nerve can be embedded in the cervical vagus in as many as half of pigs (Settell et al., 2020).

Across the cohort, stimulation location influenced the magnitude of HR responses. Multiple fiber types can drive heart rate responses. Nerve fibers responsible for cardiac changes are putatively afferent Aδ fibers from mechanoreceptors, parasympathetic efferent B fibers, and visceral afferent C fibers (Ford & McWilliam, 1986; Jones et al., 1995; Kardon et al., 1975). Consistent with studies suggesting that the HR response to VNS is primarily driven by B fibers (Chang et al., 2020; McAllen et al., 2018; Qing et al., 2018; Sabbah et al., 2011), our recorded HR responses as a function of electrode contact location show that electrodes closest to the efferent cluster of fascicles containing both motor and parasympathetic efferents elicited the most profound bradycardia. To visualize and quantify this effect, we rotated the histology to place the efferent fascicle grouping oriented toward the top of the page, as was done for the previous Aα and EMG analyses. Average maximal HR decreases were calculated and fit to a circular nerve model (Figure 6d). Visual inspection of this aggregate figure suggests that the electrode closest to the efferent cluster also evoked the most prominent HR response. HR changes across the cohort were compared between the contact closest to the efferent cluster and the contact furthest from the efferent cluster to test this hypothesis. As expected, the maximum HR decrease of the contact closest to the efferent cluster (28 ± 8 BPM) was significantly larger than the HR decrease of the contact furthest from the efferent cluster (13 ± 4 BPM) (*t*(5) = 3.6, p = 0.015, paired t-test).

### Computational Modeling

#### Quantitative Match of Modeled and In Vivo Thresholds

We implemented models based on the specific nerve morphology and electrode locations from each pig and validated the model activation thresholds for specific fiber types against the corresponding *in vivo* data (Supplementary Material: Figure 7). We found strong agreement in onset and saturation thresholds for modeled Aα fibers (13 μm MRG) in the fascicles of the efferent mode versus Aα LIFE responses and for modeled B fiber thresholds (3 μm MRG) in the fascicles of the efferent mode versus in vivo bradycardia thresholds. Current amplitudes sufficient to activate model B fibers (i.e., ~1.5 mA), also robustly activated Aα fibers across all contact locations both in the models and *in vivo* (Figure 7b-c; Supplementary Material: Figures 9-19, odd-numbered figures, panels b-c).

**Figure 7.**
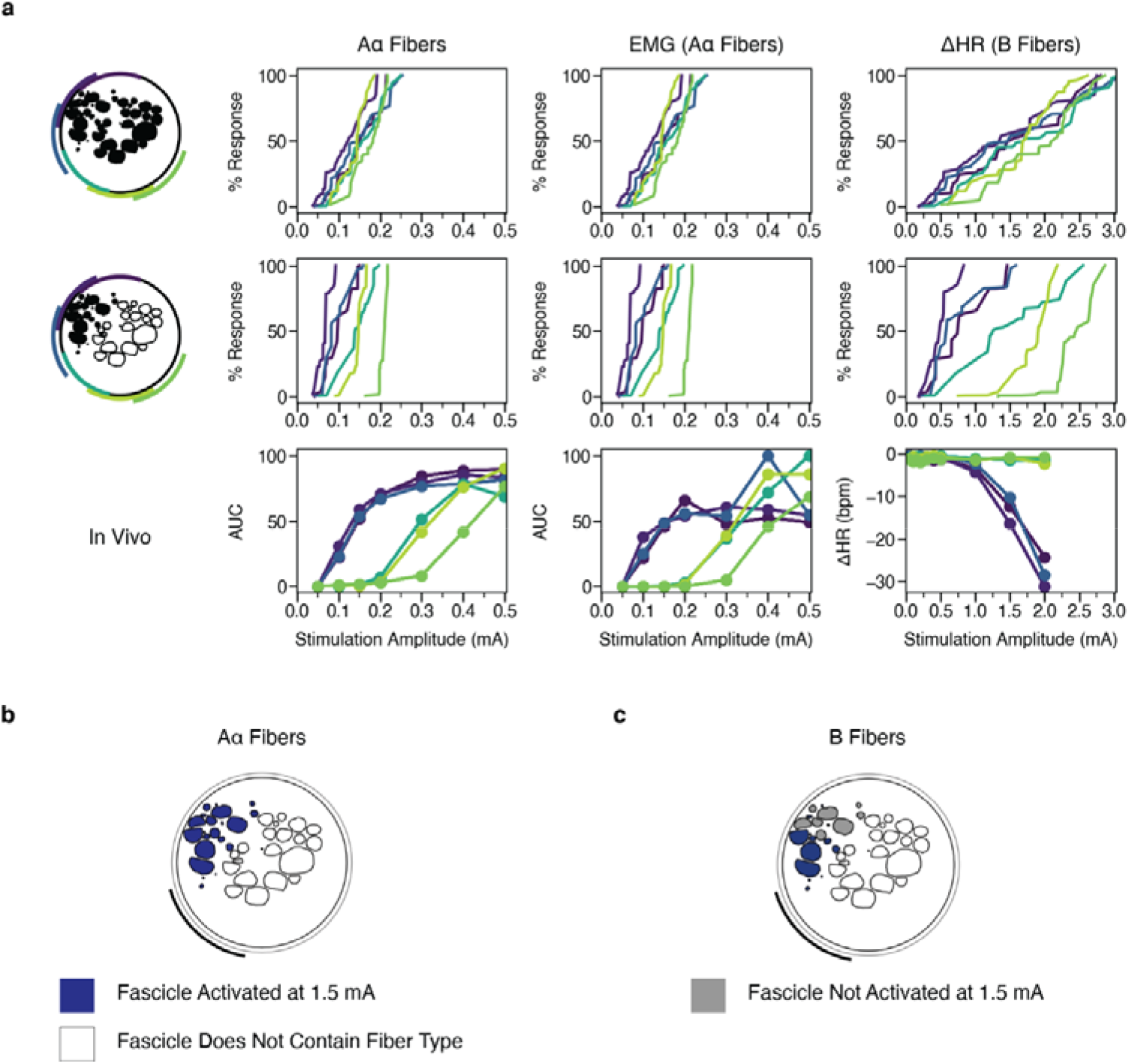
Modeling the effect of contact location on activation of Aα and B fibers without and with representation of vagotopy for Animal #2. **a,** Percent of modeled (first and second rows) and in vivo (third row) Aα and B fibers activated versus stimulation amplitude across monopolar contact location (colors). In the top row, the target fiber is positioned in the centroid of each fascicle. In the middle row, we only modeled fibers in the black fascicles (i.e., the efferent mode). **b-c,** Fascicles labeled as suprathreshold (blue) and subthreshold (gray) in response to a 1.5 mA amplitude pulse for Aα and B fibers. At this stimulation amplitude, Aα fibers are activated in all fascicles, while the B fibers are activated concomitantly only in the fascicles closest to the contact. See data for all animals in Supplementary Material: Computational Modeling Data for Additional Animals: Figures 9-19.

Models accurately predicted onset and saturation thresholds for Aα fibers (i.e., absolute model error on the same order of magnitude as the in vivo threshold resolution). Incorporating nerve vagotopy improved onset threshold accuracy for Aα fibers (i.e., 1% response: significant interaction between vagotopy and response (p = 0.038, ANOVA); p = 0.001, post hoc t-test between errors), and did not influence the errors in currents for higher activation levels (i.e., 20%, 50%, and 80% responses: p = 0.921, p = 0.263, and p = 0.487, respectively). Our models accurately predicted B fiber onset threshold magnitude as measured by onset of bradycardia. Vagotopy trended toward improved model threshold accuracy for B fiber activation (i.e., 1% response: vagotopy was almost significant in the ANOVA, p = 0.072). Models underestimated B fiber saturation thresholds (i.e., heart rate continued to decrease beyond the highest model threshold values) and did not consistently capture the contact-specific responses seen *in vivo* for the contacts furthest from the efferent mode. It should be noted that models did not include activation of Aδ or C fibers leading from baroreceptors in the aortic arch within the vagus or hitchhiking fibers from the sympathetic trunk that may also influence experimental heart rate changes, which may explain the sources of error when comparing in-vivo heart rate responses to modelled activation of parasympathetic B fibers innervating the heart.

#### Vagotopic Implications of Thresholds Across Electrode Contacts

We plotted the model dose-response curves with and without nerve vagotopy to assess whether the afferent/efferent fascicular organization determined by histology aligned with the electrode-specific ENG, EMG, and HR responses (example data provided in Figure 7; data for all animals provided in Supplementary Material: Computational Modeling Data for Additional Animals, Figures 9-19).

Without nerve vagotopy (i.e., each fiber type placed in every fascicle), we observed only small differences in the dose-response curves across contact locations for Aα and B fibers (Figure 7a: top row). Models incorporating vagotopy, with Aα and B fibers only in the fascicles of the efferent mode, improved the matches between models and experimental results (Figure 7a: middle row) and yielded more distinct contact-specific responses. Together with the experimentally documented location-specific differences in physiological responses, the model data confirm that contact location-specific responses to stimulation can be attributed to the spatially distinct locations of efferent fibers.

## Discussion

Multiple cervical VNS clinical studies report intolerable, therapy-limiting side effects resulting from neck muscle activation (De Ferrari et al., 2017; Handforth et al., 1998; The Vagus Nerve Stimulation Study Group, 1995). Biophysical principles dictate that large-diameter nerve fibers activate at lower stimulation amplitudes than small-diameter nerve fibers (Tasaki, 1953). Thus, neck muscle activation is an expected limitation since smaller diameter fibers putatively cause therapeutic effects while large diameter A fibers cause muscle contractions. However, both therapeutic effects and side effects do not have a binary on/off switch, but manifest over a dynamic range due to biophysical complexities.

The results presented here demonstrate clear location-specific differences in threshold and saturation for EMG and HR responses. Therefore, a multi-contact stimulation device such as the ImThera has the potential to shift both these curves in favorable directions, potentially increasing the effectiveness of treatment within certain limits. As the ImThera device has previously received an FDA IDE approval for human studies of stimulation of the hypoglossal nerve, the path to test these results in human studies is likely to be dramatically accelerated in comparison to completely new electrode designs.

### ImThera Cuff Design and Safety Considerations

The ImThera multi-contact cuff was approved for sale in Europe (CE Marked) to treat sleep apnea in 2021 and received an FDA Investigational Device Exemption (IDE) to conduct a U.S. Pivotal Trial in 2014. The electrode configuration was designed to stimulate the motor efferents of the human hypoglossal nerve innervating the muscles of the tongue to treat obstructive sleep apnea. As motor nerve fibers can be activated at low current amplitudes, the electrodes did not need to be designed to support higher current amplitudes needed to activate the Aδ, B, and C fibers putatively associated with the therapeutic effects of vagus nerve stimulation. The 2 mm diameter disk electrodes have a smaller surface area than the existing LivaNova bipolar helical cuff leads, and therefore the charge density applied at the same current amplitude is larger, potentially resulting in deleterious electrochemical reactions.

We followed the protocol outlined by Wilks/Ludwig et al. (Wilks et al., 2017) that was previously used to support FDA IDEs to assess the maximum safe limit for stimulation from these electrodes (Supplementary Material: Figure 1). Voltage transient analysis suggested that the average safe limitation was 5.6 ± 0.34 mA. Consequently, the maximum current applied in this acute study was 5 mA. Given that thresholds tend to increase in chronic implantations due to the formation of a scar around the electrodes, it may be necessary to pursue electrode coatings strategies to increase the electrochemical surface area of the electrodes to activate B and C fibers safely in human subjects, especially given subject-to-subject variability in the human anatomy.

It is also worth noting that each of the six ImThera contacts (contact diameter ~2 mm) spans approximately one fifth of the circumference of the pig and human cervical vagus (Hammer et al., 2018; Pelot et al., 2020). This large stimulating surface area results in overlapping coverage of the underlying neuroanatomy. Thus, it may be necessary to pursue smaller electrodes to optimize for spatial selectivity based on electrode location. However, as noted above, this would further increase charge density at a given amplitude, potentially necessitating coating strategies to safely activate higher threshold therapeutic fibers.

### Implications of Pertinent Anatomy

#### Current Escape Activates the Superior Laryngeal Branch

In two of six animals, a secondary plateau was noted in the EMG DRCs as stimulation amplitudes were increased. This phenomenon was unlikely influenced by time or repeated stimuli because stimulation amplitudes were randomized. Additionally, previous VNS work in pigs using the LivaNova helical cuff showed that high stimulation currents can indirectly activate the nearby SL branch via current escape from the insulation, leading to additional activation neck muscles in the majority of animals (Nicolai et al., 2020). The Nicolai study of the LivaNova lead used bipolar stimulation, which generates a much sharper falloff of the electrode field compared to the monopolar stimulation used in this study (1/r^2^ vs. 1/r). Nonetheless, the Nicolai data showed off-target activation of the nearby SL through current leakage from the insulation in 9 of 12 pigs at current amplitudes ranging from 0.5 to 2.5 mA.

In contrast, the ImThera cuff may be better suited for preventing current escape than a helical device in part due to a greater amount of insulation: both in terms of length from stimulating contact to the edge of insulation and thickness of insulation. This may explain the relatively limited number of subjects (2/6 herein vs. 9/12 in Nicolai) in which indirect activation was observed compared to studies using the helical design (Nicolai et al., 2020). The relative proximity of the specific contacts that resulted in off-target activation of the SL played a role within both these subjects; previous studies showed that the main branch point of the SL off the cervical vagus trunk is much closer to the accessible surgical window in pigs than in humans (Settell et al., 2020). However, the external branch of the SL still lies relatively close to the most common electrode placement in humans, and therefore somatic nerve activation via field escape from the insulation is likely still an issue in human subjects (Kierner et al., 1998).

#### Nerve Vagotopy Dictates Function

The ideal VNS intervention is one where therapy-inducing effects are maximized while concomitant therapy-impeding side effects are minimized. In this study, we observed varied location of the efferent cluster within the VN, as well as a varied relational positioning of stimulating contacts to the efferent cluster (Supplementary Material: Figure 4). Yet, we observed neck muscles and HR responses dependent on their proximity to underlying neuroanatomy. However, it is essential to note that at no contact location in any animal were thresholds for desired cardiac responses lower than the thresholds for unwanted neck muscle activation via the RL. It was, however, possible to elicit cardiac responses without causing activation of the SL through current leakage outside the insulation in four out of six animals. Further, in one of the animals experiencing muscle activation via current leakage, it was still possible to stimulate at high amplitudes in three of six contacts distant from the SL without causing additional muscle activation.

Although EMG saturation thresholds in this study were lower than thresholds for cardiac effects (Figure 8), it may be possible to habituate and desensitize the patient to these side effects (Ardell et al., 2017; Penry & Dean, 1990), to alter stimulation paradigms to favorably target a specific type of nerve fiber (Fitchett et al., 2021; Vuckovic et al., 2008), and/or to use location in conjunction with more fiber-selective stimulation strategies. Habituation and desensitization may provide enough of a shift to further increase the effectiveness of VNS due to location-specific stimulation.

**Figure 8.**
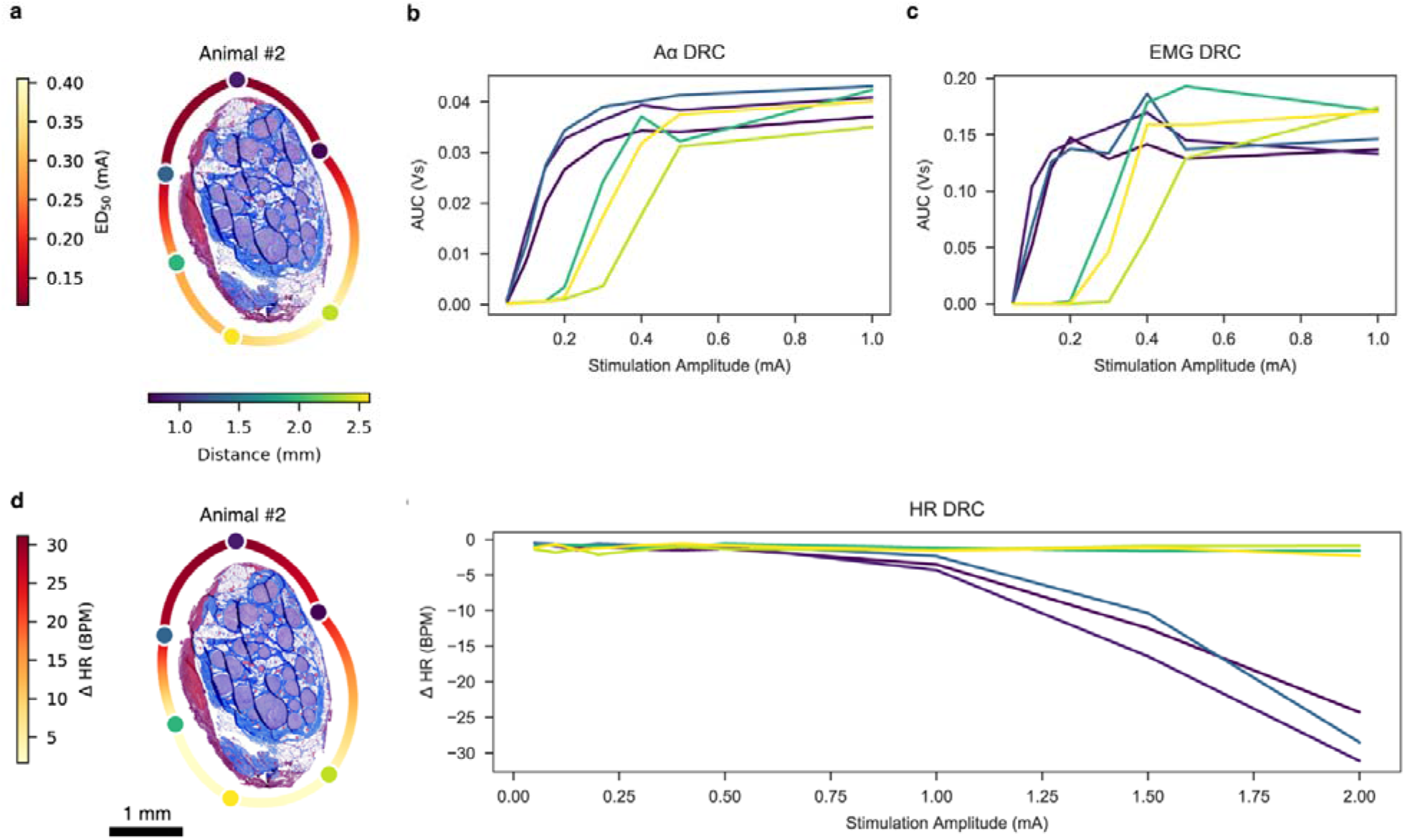
Summary figure depicting ENG, EMG, and HR DRCs and heatmaps for Animal #2. **a,** Histology of Animal #2 at the level of the stimulating cuff, rotated such that the efferent cluster is facing the top of the page. Stimulating contacts are superimposed to replicate the *in situ* condition and are characterized in color by their distance to the centroid of the efferent cluster. A heatmap was extrapolated around the perimeter of the histology to indicate changes in ED_50_ based on location. **b**, ENG dose-response curve. **c**, EMG dose-response curve. **d**, Same histology of Animal #2 as panel a, but a heatmap was extrapolated around the perimeter of the histology to indicate changes in HR based on location. **e,** HR dose-response curve.

The most striking example of contact-specific stimulation yielding varying results was evoking both bradycardia and tachycardia in Animal #1. In this animal, the two stimulating contacts closest to a hitchhiking, third grouping of fascicles traced back to the SCG of the sympathetic trunk resulted in a HR increase of over 50 BPM, while three other contacts resulted in an average decrease of 13 BPM (Figure 6, Animal #1). Cross-connections between the VN and the sympathetic trunk (Settell et al., 2020) and sympathetic fibers within the VN (Seki et al., 2014) have previously been reported. These hitchhiking fibers may be an important consideration for understanding the underlying neuroanatomy responsible for mediating competing increases and decreases in sympathetic tone.

Prior studies using the LivaNova helical cuff in canines recorded tachycardia at low current amplitudes putatively due to the activation of afferents which result in parasympathetic inhibition, with bradycardia only being induced at higher levels of stimulation (Ardell et al., 2017). This change between tachycardia at low levels of stimulation and bradycardia at high levels of stimulation is known as the ‘neural fulcrum.’ Tachycardia was not observed at low levels of stimulation in most animals in this study. However, isoflurane was used in this study which is known to act on glutamate receptors. Although blocking glutamate-mediated responses would not blunt cholinergic parasympathetic efferent to the heart, it may blunt the afferent pathway driving reflexive parasympathetic inhibition. Another possibility is that the relatively simple cervical vagus fascicular organization in canines consisting of 1-3 fascicles (Onkka et al., 2013; Yoo et al., 2013) has significantly different functional responses to VNS than the more complex fascicular organization of the pig vagus (Pelot et al., 2020; Settell et al., 2020).

### Computational Modeling

Our modeling results support the main takeaway from the in vivo data: the ImThera cuff in monopolar configuration provides no opportunity to activate B fibers in the pig vagus nerve before robust concomitant activation of side effect producing Aα fibers. Furthermore, our paired experimental and modeling data highlight the importance of accounting for individual-specific nerve morphology, nerve vagotopy, and distinct fiber types when attempting to target specific physiological responses.

Although previous studies modeled stimulation of individual-specific nerves to target selectively groups of fascicles that innervate distinct muscles (Schiefer et al., 2008; Brill et al., 2011; Kent et al., 2013), we are the first to quantify selective stimulation of functionally distinct fascicles in the vagus nerve with individual-specific models. Models’ accuracy was critically enabled by incorporating controlled cuff orientation, nerve vagotopy and accurate conduction velocities for the fibers driving distinct physiological responses, and models that correspond one-to-one with nerves stimulated in vivo achieved the strongest contact specific match for Aα and B fiber thresholds.

Models enable rapid, reproducible, and efficient quantification of therapy design choices on nerve response to stimulation. Specifically for VNS, patients need reduced side effects, which will require avoiding activation of side effect producing fibers both within the nerve trunk proper and in nearby branches (e.g., the SL) from current escape (Nicolai et al., 2020).

### Limitations

Cardiac effects of isoflurane and fentanyl are well characterized, but their competing interaction is complex. Isoflurane can lead to vagal inhibition, causing an increase in HR (Huang et al., 1997; Kato et al., 1992; Picker et al., 2001). In contrast, fentanyl decreases HR due to an increase in vagal tone (Griffioen et al., 2004; Laubie et al., 1977). All recordings analyzed during this study were obtained during periods when isoflurane concentration was less than 2%. Efforts were made to keep the isoflurane concentrations minimal during the entire procedure. However, due to animal safety and welfare concerns, isoflurane concentrations occasionally rose above 2%. Synaptic blunting caused by isoflurane anesthesia may require activation of a larger number of efferent parasympathetic fibers or afferent fibers from mechanoreceptors to elicit a HR response, and therefore the anesthesia may have affected the activation and saturation thresholds, as well as the total change in HR (Baumgart et al., 2015; Herring et al., 2009).

Putative nerve fibers responsible for cardiac changes are afferent Aδ fibers from mechanoreceptors, parasympathetic efferent B fibers, and visceral afferent C fibers (Ford & McWilliam, 1986; Jones et al., 1995; Kardon et al., 1975). However, we were unable to obtain LIFE recordings from Aδ, B, or C fibers. Prior studies suggest that C fibers are not activated within clinically tolerated stimulation intensities (Yoo et al., 2013) and that Aδ fibers may only play a small role in the cardiac response, especially in the right VN (Ito & Scher, 1973). Therefore, B fiber measurements are the best predictors for cardiac activity (Qing et al., 2018; Sabbah et al., 2011). However, we were unable to record B fiber activity and analyze this link in the pathway between applied stimulus and cardiac response. This is unsurprising, as other studies have either only intermittently captured recordings from these fiber types (Nicolai et al., 2020), or demonstrated that measuring changes in potential due to these fibers may require surgically removing the epineurium of the VN in large animal models (Yoo et al., 2013). Instead, we recorded changes in HR as therapeutic effect and recorded EMG activity as side effects, analogous to clinical neck pain, throat tightening, cough, dyspnea, or hoarseness (De Ferrari et al., 2017; Handforth et al., 1998; The Vagus Nerve Stimulation Study Group, 1995), although these may be oversimplifications. We may have therefore come to ambitiously broad conclusions, albeit having sound inferential logic, meriting further validation in future studies.

Acute surgical trauma often resulted in fluid and edema at or near the electrode/nerve interface. Such fluid was wicked away via surgical gauze as needed. This fluid buildup may have impacted certain variables such as the spatial distribution of applied currents and thus stimulation thresholds. In addition, the surgery may have impacted additional biophysiological phenomena, such as indirect activation of nearby anatomical structures. Therefore, further studies with chronically implanted electrodes are needed to understand the roles that edema, fluid, scar tissue formation, and biofouling play in the responses described in this paper.

The computational models have limitations to their complexity for maintaining feasibility. Specifically, we assumed a constant nerve cross section along the model length, a radially uniform correction to account for tissue shrinkage during histology, and a physics-based method for reshaping the nerve to a circular cross section in the cuff electrode and repositioning the fascicles accordingly. Further, we did not model distributions of diameters of each fiber type, which could allow some of the largest B fibers to be recruited before the smallest Aα fibers.

There were limitations in the validation of model data against corresponding in vivo measurements. The in vivo threshold data that we used to compare with the modeling data may lack sufficient signal-to-noise to detect the onset and saturation current amplitudes for specific nerve fiber types, especially small diameter nerve fibers, which generate a comparatively weaker extracellular signal. In the models, the lowest and highest thresholds across fascicles are known, but we do not know precisely which fascicles contributed to the associated ENG/EMG signal. Further, we were limited in the resolution of amplitudes tested in vivo; therefore, for each onset and saturation threshold in the in vivo data, there is an uncertainty range within which the response started or saturated. We assumed that the Aα and B fibers were both uniformly distributed across the fascicles of the efferent mode, but they may each be grouped within those fascicles, which could provide additional selectivity control.

## Conclusion

Therapeutic outcomes of VNS are limited in part by the stimulation-induced side effect of neck muscle activation. In this study, we sought to characterize the fundamental relationship between the underlying anatomical organization of fiber types within the cervical vagus trunk in pigs and the functional outcomes of location-specific electrical stimulation in terms of both effect and side effect. Functional and histological data were used to develop and validate computational models of location-specific activation. Both *in vivo* and *in silico* results demonstrate location-specific differences in threshold and saturation for responses related to effect and side effect. Therefore, a multi-contact stimulation device such as the ImThera cuff may exploit the underlying fascicular bimodality of efferent versus afferent fiber types and shift the ratio of therapeutic effect to noxious side effect in a favorable direction, potentially increasing the effectiveness of treatment.

Previous studies determined the importance of certain VNS parameters such as stimulation amplitude and frequency. Here, we have demonstrated an equally important parameter: stimulation location.

## Acknowledgments

Funding from NIH SPARC Program OT2 OD025340 and LivaNova USA Inc.

Thanks to Jared Ness and Ian Baumgart for their help in collecting electrophysiological data and to Daniel Marshall for segmenting the nerve histology used to define nerve cross sections in ASCENT. A sincere thank you to Nishant Verma and Dr. Wolf-Ekkehard Blanz for proofreading the article.

The opinions expressed in this article are the author’s own and do not reflect the view of the National Institutes of Health, the Department of Health and Human Services, or the United States government.

## Footnotes

1 In previous studies, we conducted immunohistochemistry with an antibody against choline acetyltransferase on the pig cervical vagus nerve. We demonstrated that cholinergic fibers consistent with both motor and parasympathetic efferents are preferentially clustered in one of the two bi-modal groupings of fascicles (Settell et al., 2020).

2 The most visible error bars can be seen when closely examining Animal #4’s Aα DRCs. The narrowness of the error bars is attributed to the fact that over the 750 pulses per train, the evoked neural signal from pulse to pulse is remarkably consistent.

