## Supplementary Material for "Spatially selective stimulation of the pig vagus nerve to modulate target effect versus side effect"

### Supplementary Information

#### Methods

##### Electrochemical Testing and Validation

To determine the suitability and safe stimulation limits for the ImThera device, we conducted electrochemical impedance spectroscopy (EIS), voltage transient (VT) analysis, and cyclic voltammetry (CV) as previously published by our group for investigating novel devices and for FDA IDE submissions (Trevathan et al., 2019; Wilks et al., 2017). EIS was performed to determine uniformity of measurements across the six stimulating contacts (Figure 1a) and to compare the resultant Bode plots to those of platinum-iridium alloys established in literature (Petrossians et al., 2011).

To establish safe stimulation limits, we conducted VT and CV studies. VT studies determine which stimulation currents do not exceed water oxidation potential limits based on the loci of the most negative and most positive polarizations (E_mc_ and E_ma_, respectively) recorded per stimulation channel (Cogan, 2008). For platinum-iridium electrodes, E_mc_ must be greater than -0.6 V and E_ma_ must be less than 0.8 V. These safety limits were not exceeded at or below stimulation currents of 5 mA (Figure 1c). Based on these excursions across all stimulation contacts, a maximum stimulation current of 5.6 ± 0.34 mA (mean ± SD) was calculated. From CV measurements, a cathodic capacitive storage charge (CSC_c_) of 18 mC cm^-2^ ± 0.43 mC cm^-2^ was calculated. This value is well above the charge injection limits for platinum-iridium alloys of 0.05 mC cm^-2^ – 0.15 mC cm^-2^ (Cogan, 2008; Rose & Robblee, 1990). Additionally, these charge injection limits for platinum-iridium alloys (0.05 mC cm^-2^ – 0.15 mC cm^-2^) were used to calculate maximum stimulation currents. These calculations yielded maximum stimulation currents of 5.5 mA (at 0.05 mC cm^-2^) and 17 mA (at 0.15 mC cm^-2^).

These data indicate that the stimulation contacts within the ImThera device have electrochemical properties suitable for neural stimulation and/or recording.

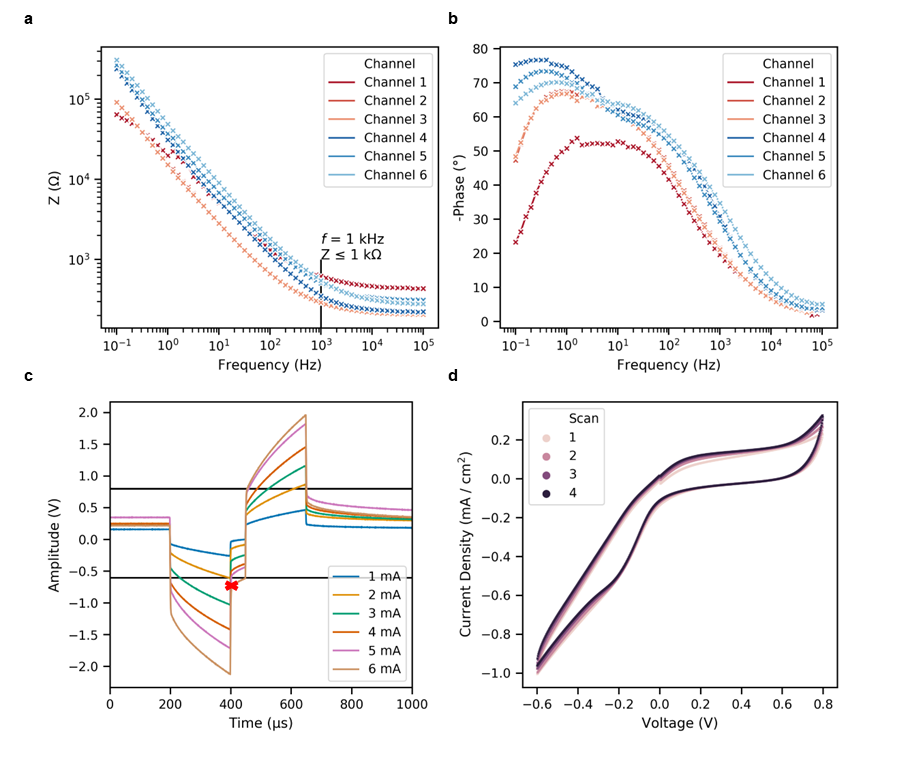

**Figure 1. a.** Magnitude of the impedance (Z) versus frequency for all channels of the ImThera device. **b.** Plot of the negative of the phase of the impedance versus frequency. An inverse parabolic shape centered between 0 Hz and 10 Hz was characteristic across stimulation channels. **c.** VT recordings from a representative channel. Stimulation currents higher than 3 mA caused the E_mc_ to measure below -0.6 V. The red ‘X’ indicates the first E_mc_ below -0.6 V (from 3 mA stimulation). **d.** CVs at a steady state from a representative channel. CVs reveal electrochemical reactions at the electrode. Additionally, the CSC_c_ can be determined by calculating the time integral of the cathodic current.

##### Stimulation Waveform

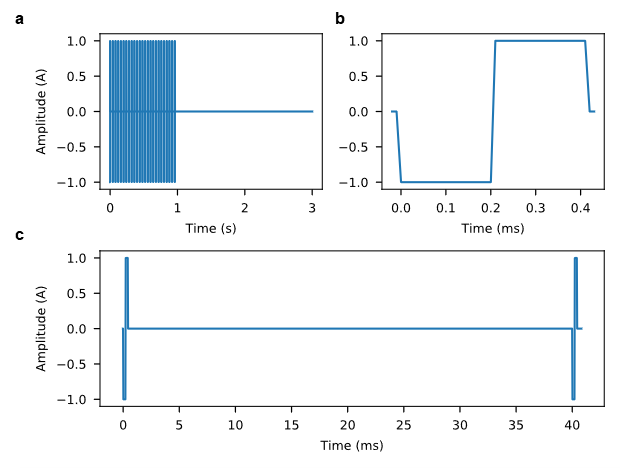

**Figure 2. a.** Pulse train of 25 pulses at 25 Hz. **b.** Each pulse is a charge-balanced, biphasic rectangular pulse with 0.2 ms duration per phase and 0 ms interphase delay for a total pulse duration of 0.4 ms. **C.** Snapshot of two pulses within the pulse train to show delay between pulses at 25 Hz (40 ms). Note the “intact” condition consisted of 750 pulses at 25 Hz, rather than the 25 pulses shown here for illustrative purposes.

##### Nerve Fiber Type Overview

**Table 1**. Generalized overview of fiber types and their pertinent functions, attributes, and implications for VNS (Manzano et al., 2008).

| **Fiber Type** | **General Function** | **Putative Target of VNS? (Yes/No)** | **Conduction Velocity (m/s)** | **Fiber Diameter (μm)** |
| --- | --- | --- | --- | --- |
| A-α | Motor efferent | N – avoid; leads to side effect | 70-120 | 12-22 |
| A-β | Somatic afferent | N | 30-70 | 5-12 |
| A-γ | Motor efferent | N – avoid; leads to side effect | 15-30 | 2-8 |
| A-δ | Visceral & somatic sensory afferent | Y (visceral afferent from baroreceptors) | 5-30 | 1-5 |
| B | Parasympathetic efferent | Y | 3–15 | ≤ 3 |
| C | Visceral sensory afferent | Y | 0.6–2.0 | 0.1-1.3 |

##### Evoked Compound Action Potential Calculation via “Peaks” Methodology

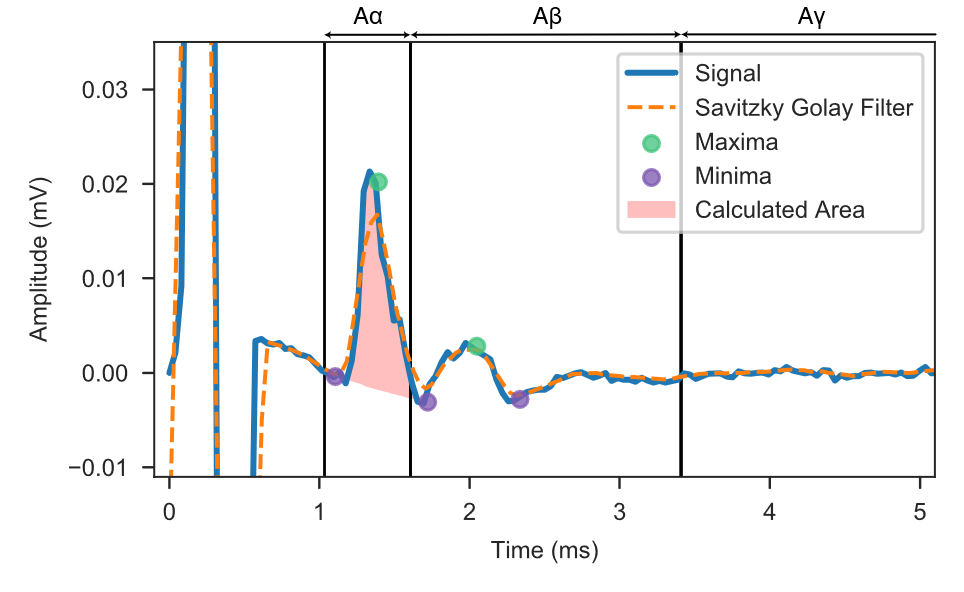

**Figure 3**. The raw signal (blue) was filtered using a Savitzky-Golay Filter. The filtered signal (dashed orange) was then analyzed for the local maxima (green circles) within each time window, i.e., for each fiber type. The closest minima before and after each local maximum (purple circles) were found via second derivative and then defined as the bounds of each eCAP component. The definite integral of a straight line between these points was calculated and subtracted from the definite integral of the raw signal, resulting in the area shaded in red. Various factors could affect the characteristics of the decay of the stimulation artifact (e.g., fluid in the surgical pocket, distance to the electrical ground, amplitude of stimulation); therefore, we used this analysis approach rather than subtracting a potentially mismatching template artifact.

##### Interpolation and Statistics

In the “intact” condition, the pulse train (per unique stimulation amplitude) consisted of 750 pulses (similar to **Figure 2**). In the time between these individual pulses (epoch), ENG and EMG activity was recorded. This resulted in 750 epochs with four ENG recordings and two EMG recordings (four EMG recoding channels, two EMG recording channels). We sampled with replacement the epochs within each pulse train 750 times. We took the mean of the data from the sampled epochs, resulting in four average ENG (one per channel) and two average EMG (one per channel) data sets per pulse train. From these averages, the AUC or V_RMS_ was calculated (resulting in four values per ENG and two values per EMG). These values were then averaged, resulting in one AUC per ENG and one AUC per EMG. This entire process was repeated 1,000 times to establish a confidence interval.

##### Histology, Stimulating Contacts, and Rotation

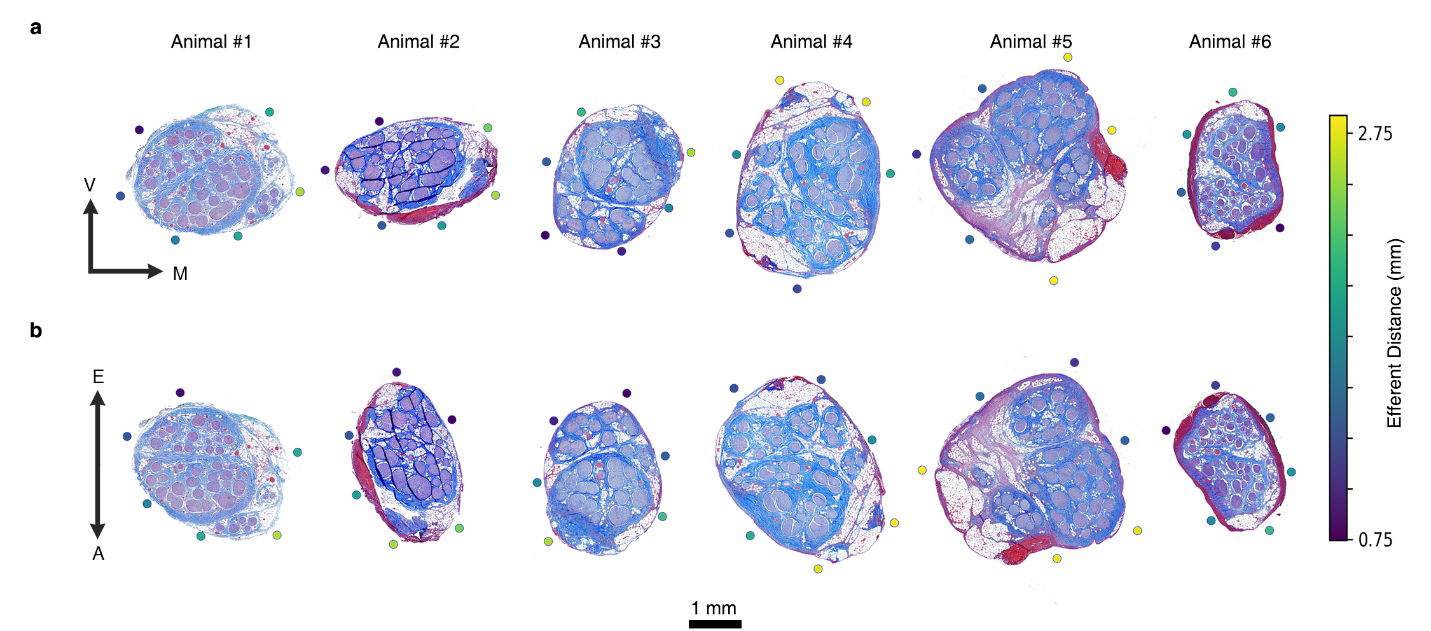

**Figure 4. a.** *In Situ* depiction of nerve histology across the cohort with stimulating contacts superimposed, rotated to align the ventral (V) and medial (M) directions. The distance of each contact to the centroid of the efferent cluster was calculated and depicted by color. **b.** Contact/histology rotation such that the efferent (E) cluster is pointing towards the top of the page and the afferent (A) cluster is pointing towards the bottom of the page.

##### Computational Modeling

For each pig, we segmented the epineurium and fascicles in the histological cross section at the level of the ImThera cuff using Nikon’s NIS-Elements Ar software (v5.02.01, Build 1270, Nikon Instruments Inc.) as described previously (Pelot et al., 2020) (Figure 5). We scaled each nerve cross section radially by 16.67% to correct for tissue shrinkage during histological processing (Boyd & Kalu, 1979). Using ASCENT’s physics-based fascicle repositioning algorithm, we deformed the outer nerve boundary to be circular while maintaining its cross-sectional area and ensuring that the fascicles were not overlapping after deformation. The TIFs of each segmented nerve morphology and the JSON configuration files used are in Supplementary Material: ascent_inputs.zip.

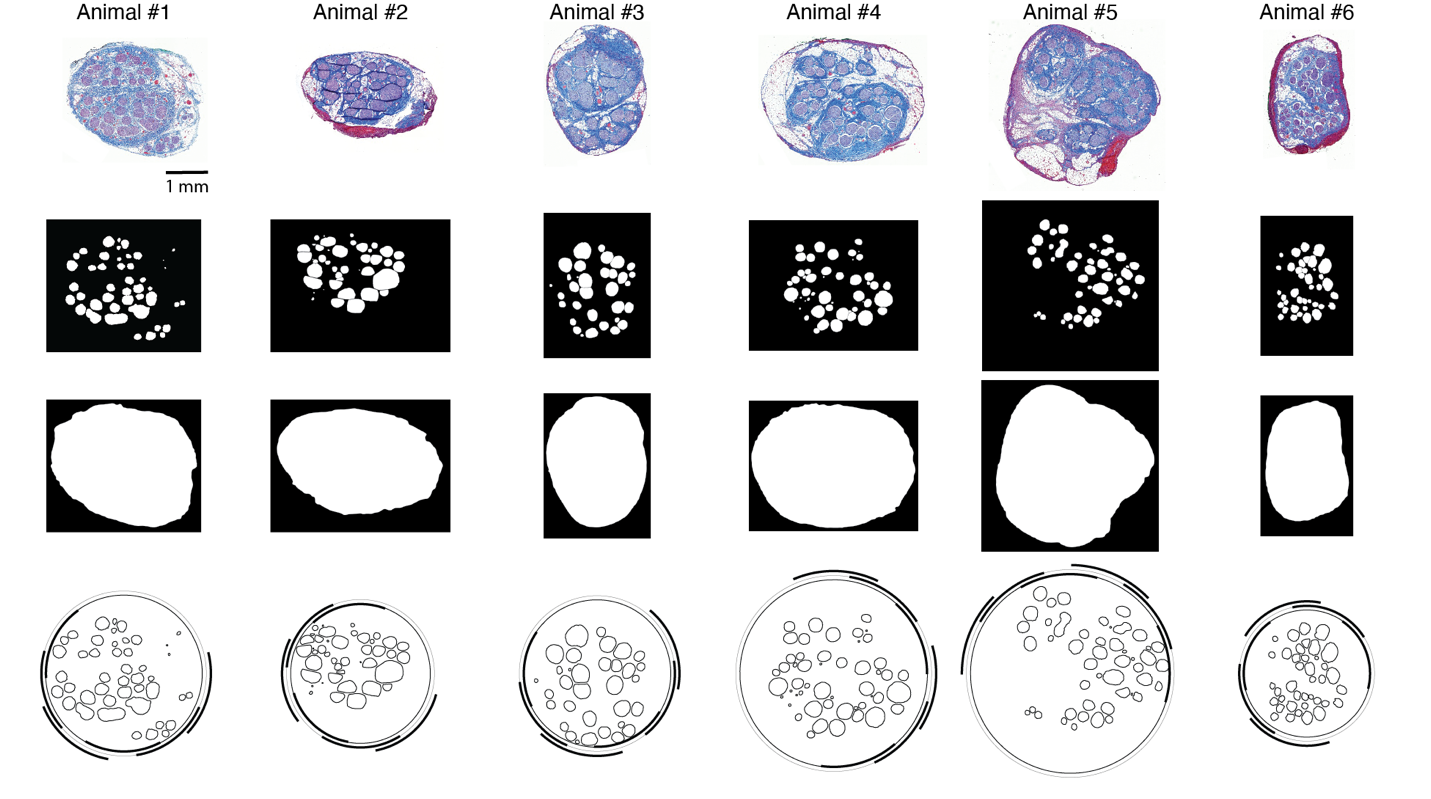

Figure 5. Animal-specific nerve morphologies modeled in ASCENT using segmented histology from nerves stimulated in vivo. First row: Nerve histology for six pigs. Second row: Segmented endoneurium (i.e., INNERS). Third row: Segmented epineurium. Fourth row: Fascicle and nerve traces after tissue shrinkage correction and ASCENT’s physics-based nerve deformation to the circular inner diameter of the ImThera (six contact locations shown).

We used the ASCENT (“Automated Simulations to Characterize Electrical Nerve Thresholds”) pipeline v1.1.0 (Musselman et al., 2021) to model VNS for all six pigs with the ImThera cuff electrode using individual-specific nerve morphology from histology. ASCENT built, meshed, and solved finite element models (FEMs) using COMSOL Multiphysics v5.4 (Burlington, MA). We extruded the nerve cross section 25 mm in the longitudinal dimension to define the three-dimensional nerve (Figure 6). We ensheathed each fascicle with perineurium, modeled as a sheet resistance (i.e., “contact impedance” in COMSOL) with values determined by multiplying the material resistivity by the perineurium thickness (Pelot et al., 2019; Weerasuriya et al., 1984); perineurium thickness was calculated for each fascicle using our previously published linear relationship between fascicle diameter and perineurium thickness for the pig vagus nerve (Pelot et al., 2020). Using ASCENT’s library of part primitives for assembling custom cuff electrodes, we implemented the ImThera six-contact cuff electrode (Figure 6), which was stretched open from its resting diameter of 3 mm if needed to accommodate the nerve. We placed the cuff halfway along the nerve length (25 mm), and we used tissue dye visible in the histology as a landmark to match cuff rotation in the model with the in vivo placement. The preset cuff JSON file used is in Supplementary Material: ascent_inputs.zip. We modeled a 100 μm layer of saline over all surfaces of the cuff. We modeled the surrounding tissue as muscle (25 mm length, 10 mm diameter).

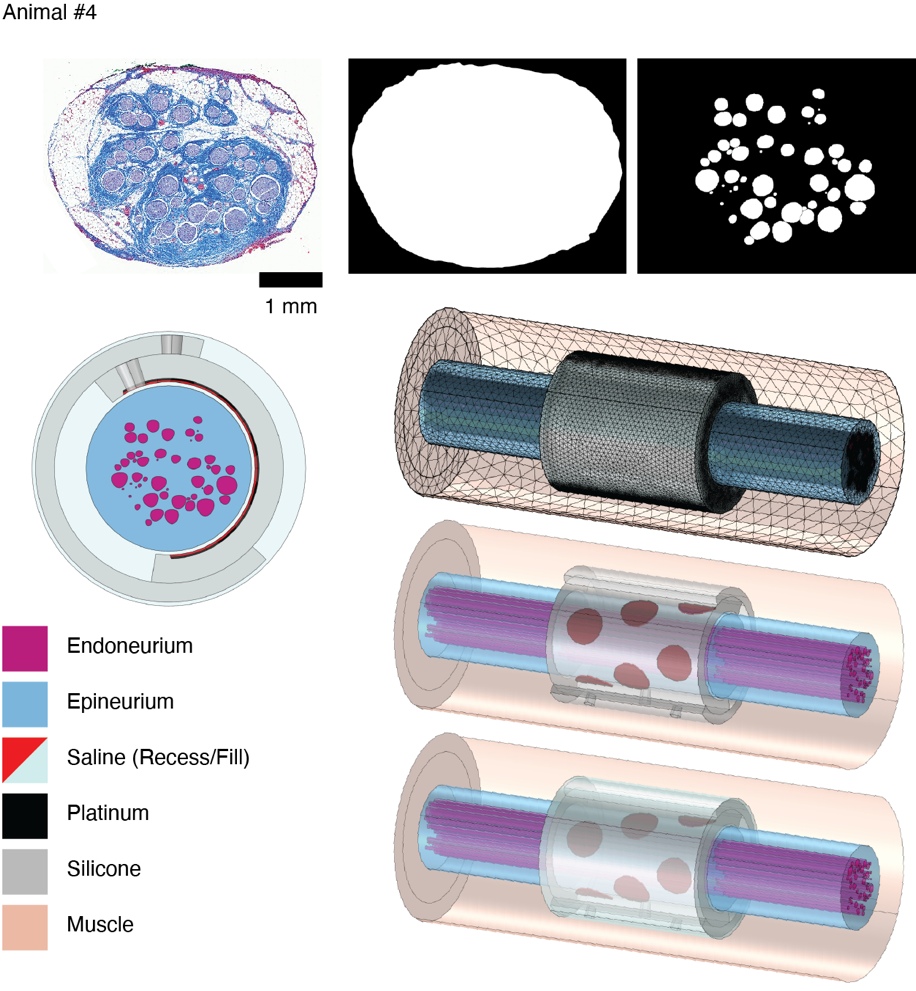

**Figure 6.** Example finite element model of an animal-specific pig vagus nerve (Animal #4) and ImThera cuff built using the ASCENT pipeline to replicate the in vivo electrophysiology experiment. Each fascicle was sheathed in perineurium (not shown) which we modeled with a thin layer approximation. The scale bar applies to the three panels in the top row: histology (left), segmented epineurium (middle), and segmented endoneurium (right).

We assigned realistic material and tissue properties to modeled domains (Table 2, Figure 6). The perineurial contact impedance for each fascicle was calculated using our published relationship between fascicle diameter and perineurium thickness for pig VN (thk_peri_ = 0.02547*d_fasc_ + 3.440 µm) (Pelot et al., 2020) and using the perineurial conductivity at 37 ^o^C, 1149 Ω-m (Pelot et al., 2019; Weerasuriya et al., 1984).

**Table 2.** Table of conductivities for the modeled tissues and materials. The endoneurium and muscle are anisotropic, with different conductivities in the x, y, z directions.

| **Material** | **Conductivity [S/m]** | **Reference** |
| --- | --- | --- |
| Endoneurium | 0.167, 0.167, 0.571 | (Pelot et al., 2019; Ranck & BeMent, 1965) |
| Epineurium | 0.159 | (Grill & Mortimer, 1994; Pelot et al., 2017; Stolinski, 1995) |
| Muscle | 0.086, 0.086, 0.35 | (Gielen et al., 1984) |
| Platinum | 9.43 x 10^6^ | (de Podesta et al., 1996) |
| Saline | 1.67 | (Horch, 2017) |
| Silicone | 1 x 10^-12^ | (Callister & Rethwisch, 2012) |

We meshed the FEMs with 17,207,060 to 24,962,721 tetrahedral elements. ASCENT solved Laplace’s equation for each FEM with 1 mA delivered to each contact independently (i.e., solved six times) and the outer boundaries of the model grounded to serve as a distant return. For each of the solved COMSOL models, ASCENT sampled the resulting electric potentials along the length of the nerve at the centroid of each fascicle. The sampled potentials were applied to biophysically-realistic mammalian myelinated model fibers (McIntyre-Richardson-Grill model (MRG); (McIntyre et al., 2002, 2004; Musselman et al., 2021)) as a time-varying waveform using NEURON v7.6 (Hines & Carnevale, 1997). We centered the middle node of Ranvier at half of the nerve length in the FEM in all simulations. We modeled fiber diameters that correspond to Aα and B fiber conduction velocities. Specifically, for Aα fibers, we modeled fiber diameters with conduction speeds within the range of experimentally recorded LIFE signals (i.e., Aα: MRG 13 µm, 73.2 m/s (70-77 m/s)); while a larger range of conduction speeds was considered in analyzing the in vivo data, this range capture the peaks of the detected signals in the in vivo recordings. For B fibers, we modeled 3 µm diameter MRG fibers, consistent with published distributions of fiber diameters (Fazan & Lachat, 1997; Licursi de Alcântara et al., 2008; Mei et al., 1980; Soltanpour & Santer, 1996).

We used a binary search algorithm (1% resolution) to identify the activation threshold for each fiber in response to a symmetric biphasic pulse (200 µs/phase, as used in vivo). We delivered the pulse at t = 1 ms and recorded the transmembrane potential at the node of Ranvier closest to 90% of the fiber length during the 50 ms simulation. We used a time step of 1 µs and backward Euler integration.

We conducted convergence studies of the activation thresholds to ensure that the model length, model radius, and mesh resolution were sufficient to ensure numerical accuracy and to avoid edge effects (Howell & Grill, 2014). Using a binary search algorithm, we computed activation thresholds for 2 µm diameter mammalian myelinated nerve fibers (McIntyre et al., 2002) in response to a 100 µs monophasic pulse at ~20 randomly chosen fiber locations in each fascicle. This study used a different model of a pig VN and monopolar cuff electrode, but we expect the parameters to be suitable across individuals and similar cuff electrodes.

First, with the radius of the model held constant at 10 mm and using a very fine mesh, we reduced the length of the model from 100 mm (the “largest” model in the proceeding text) by half until there was a change in activation thresholds greater than 2% compared to the largest model, resulting in a length of 25 mm. Second, with the length of the model held constant at 25 mm and using the same very fine mesh, we reduced the radius of the model from 10 mm by half until there was a change in thresholds greater than 2% compared to the largest model, resulting in a radius of 5 mm. After determining appropriate model dimensions, we reduced the mesh density of the model by half until there was a change in thresholds greater than 2% compared to the largest model. We checked that the model with converged length, radius, and mesh had thresholds within 2% of the largest model. Lastly, we rechecked the model length with larger diameter fibers (16 μm) to ensure that they had enough nodes of Ranvier; we found no change in fiber thresholds between FEMs that were 25 versus 40 mm long (i.e., 17 versus 27 nodes of Ranvier).

We compared model and in vivo Aα fiber thresholds across response levels. Percent model Aα fiber responses (i.e., 1, 20, 50, 80%) were calculated by adding the cross-sectional areas of all fascicles with thresholds less than or equal to incremental stimulation amplitudes and dividing the result by the sum of all fascicle cross-sectional areas. In vivo Aα thresholds were determined from the Hill model equation fits of the dose-response curves, from which we derived the stimulation amplitudes at which 1, 2, 50, and 80% of the maximum response occurred for a given contact.

We also compared model and in vivo B fiber onset thresholds. Model B fiber onset thresholds were defined as the minimum threshold across all fascicles containing B fibers for a given contact. From the in vivo heart rate data, we defined B fiber onset threshold as the stimulation amplitude at which the heart rate first dropped by more than 2 bpm, and then calculated the onset threshold as the midpoint between that stimulation amplitude and the next lowest amplitude tested. Contacts for which the heart rate never dropped by more than 5 bpm at any stimulation amplitude were excluded from our analyses.

For each metric (i.e., Aα or B fibers for a given response level), we calculated the model error by subtracting the model threshold from the in vivo threshold. Analyses and plotting were completed in Python v3.7 (Van Rossum & Drake, 2009).

#### Results

##### Computational Modeling Data for Additional Animals

###### Validation Summary

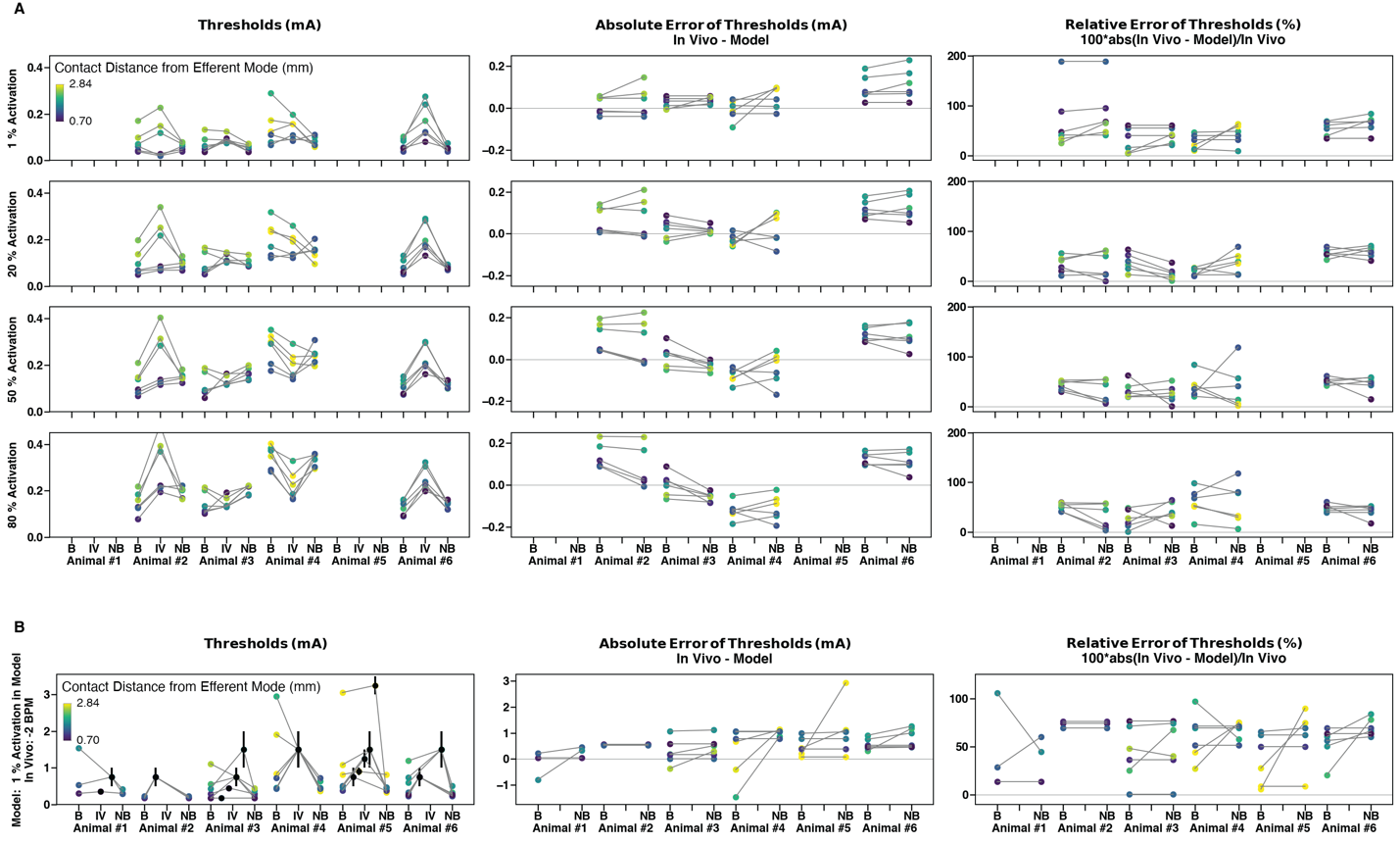

Figure 7. Comparison of modeled to in vivo thresholds for A and B fibers across animals and response levels. Each data point represents the value for a single contact location, which is color-coded by distance from the efferent mode containing the target fibers. The first, second, and third columns show raw threshold values, absolute error of thresholds, and relative error of thresholds, respectively. a, Aα fiber thresholds and errors for models with bimodal distribution of fibers (“B” on x axis; i.e., 13 µm fibers only modeled in the fascicles of the efferent mode), models without bimodal distribution of fibers (“NB” on x axis; i.e., 13 µm fibers modeled in all fascicles), and in vivo (“IV” on x axis; from Aα LIFE recordings). The rows show data for 1, 20, 50, and 80% response levels. Animals #1 and #5 did not have in vivo Aα LIFE recordings due to contamination from the stimulus artifact. b, B fiber onset thresholds and errors between modeled and in vivo thresholds. The in vivo data points averaged the lowest amplitude that elicited at least a 2 BPM change in heart rate and the next lowest amplitude; the error bars span to those nearest stimulation amplitudes. The x axis labels are the same as in panel a, except 3 µm fibers were modeled instead of 13 µm fibers.

###### Animal #1

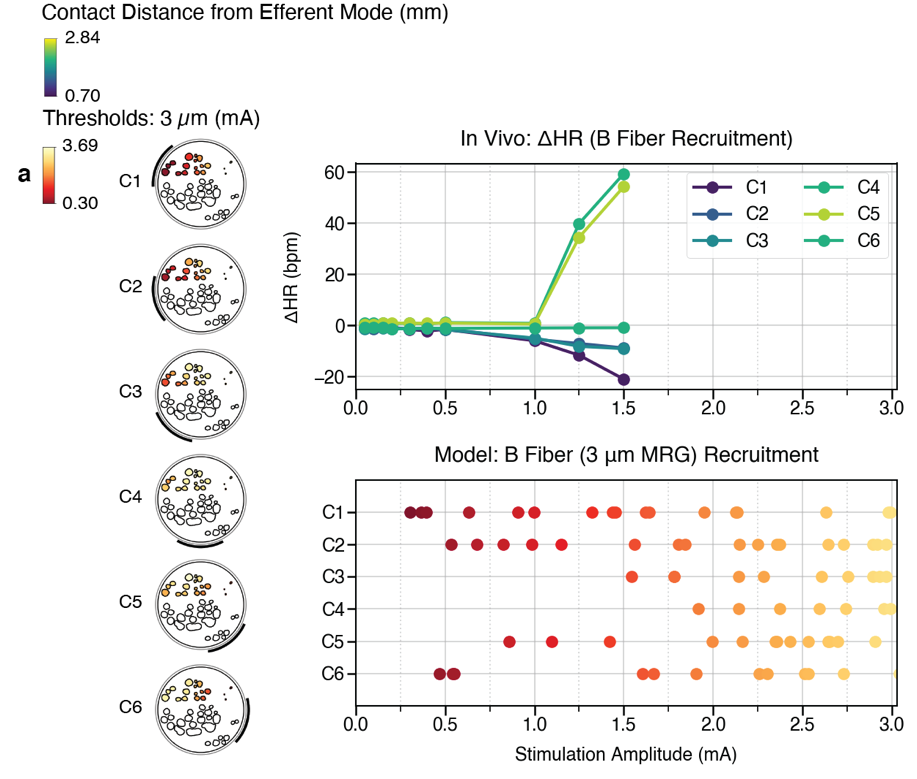

Figure 8. Comparison of in vivo thresholds to model thresholds in the pig cervical vagus nerve with the ImThera cuff electrode for Animal #1. a, B fibers modeled in the efferent mode compared to change in animal heart rate for a range of amplitudes.

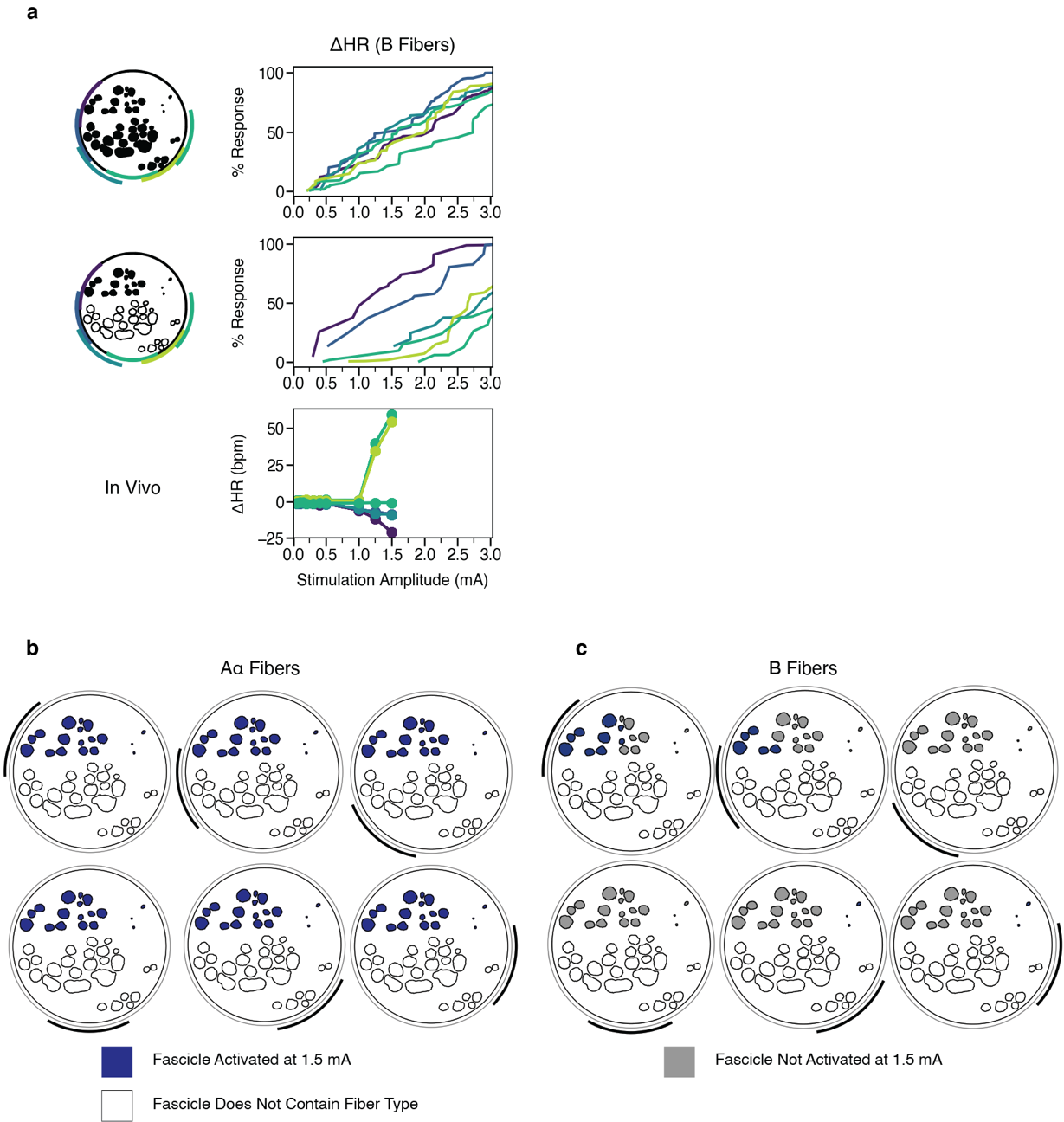

Figure 9. Animal #1 model and in vivo recruitment curves for B fibers. a, Percent of modeled (top two rows) and in vivo (bottom row) B fibers activated versus stimulation amplitude across monopolar contact configurations (colors). In the top row, we modeled a B fiber in the centroid of each fascicle. In the middle row, we only modeled fibers in the black fascicles (i.e., the efferent mode). b-c, Fascicles labeled as suprathreshold (blue) and subthreshold (gray) in response to a 1.5 mA amplitude pulse for Aα and B fibers, respectively, for Animal #1. At this stimulation amplitude, Aα fibers are activated in all fascicles, while the B fibers are activated concomitantly only in the fascicles closest to the contact.

###### Animal #2

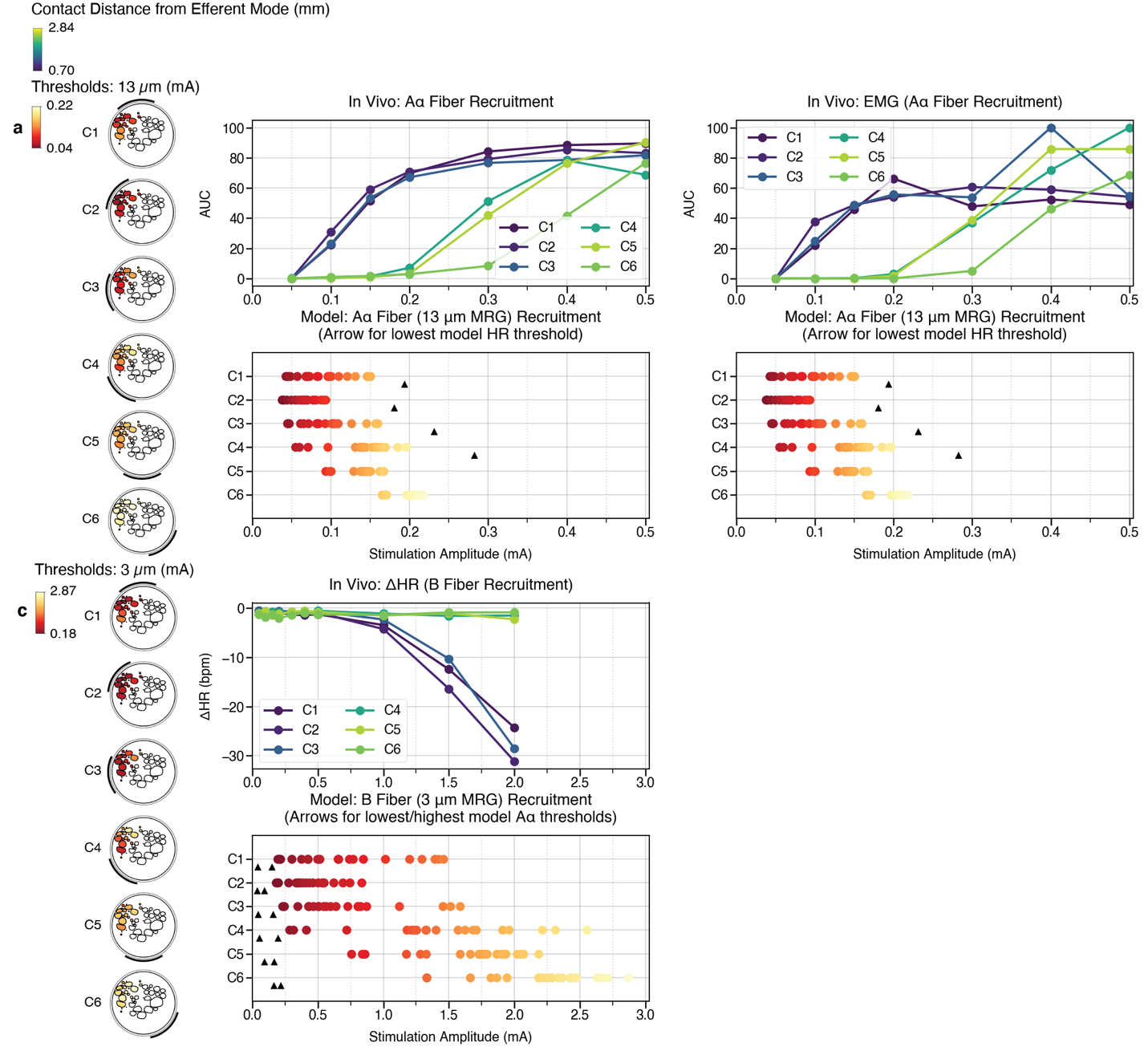

Figure 10. Comparison of in vivo thresholds to model thresholds in the pig cervical vagus nerve with the ImThera cuff electrode for Animal #2. a, Aα fiber thresholds modeled in the efferent mode compared to LIFE signals with corresponding conduction velocity. The arrows in the bottom panel indicate the lowest model B fiber threshold (i.e., heart rate onset threshold) for the corresponding contact, if the threshold fell on the x-axis range. In this case, all Aα fibers are activated before any model B fibers. b, Same modeling data as panel a plotted beside the laryngeal EMG signal. c, B fibers modeled in the efferent mode compared to change in animal heart rate for a range of amplitudes. The arrows in the bottom panel indicate the lowest and highest model Aα threshold for the corresponding contact. In this case, all Aα fibers are activated before any model B fibers.

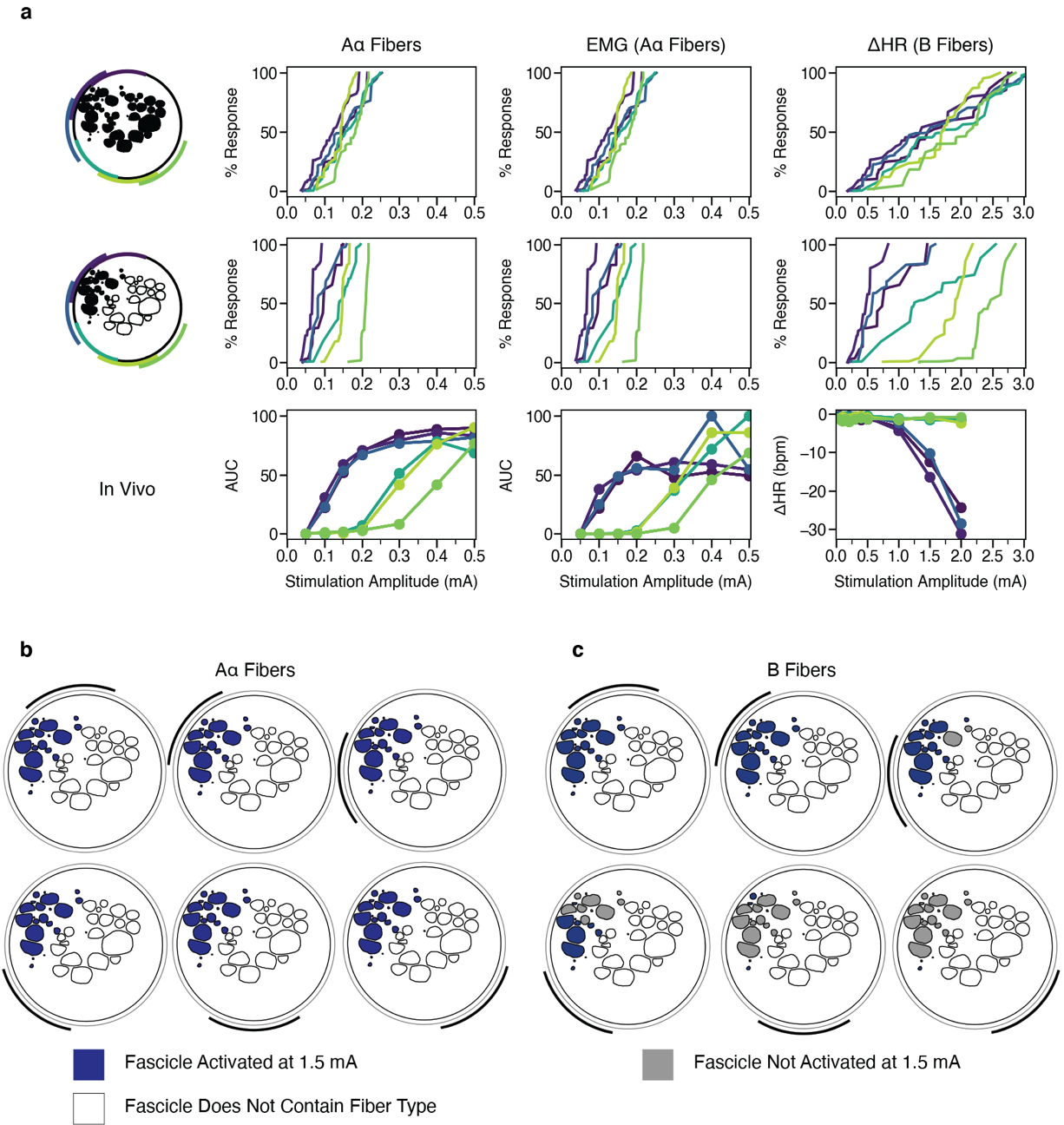

Figure 11. Animal #2 (with and without vagotopy) and in vivo recruitment curves for Aα and B fibers. a, Percent of modeled (top two rows) and in vivo (bottom row) Aα and B fibers activated versus stimulation amplitude across monopolar contact configurations (colors). b, Percent of modeled (top two rows) and in vivo (bottom row) Aβ fibers activated versus stimulation amplitude across monopolar contact configurations. In the top row of panels a and b, we modeled an Aα or B fiber in the centroid of each fascicle. In the middle row, we only modeled fibers in the black fascicles (i.e., the efferent mode). b-c, Fascicles labeled as suprathreshold (blue) and subthreshold (gray) in response to a 1.5 mA amplitude pulse for Aα and B fibers, respectively, for Animal #2. At this stimulation amplitude, Aα fibers are activated in all fascicles, while the B fibers are activated concomitantly only in the fascicles closest to the contact.

###### Animal #3

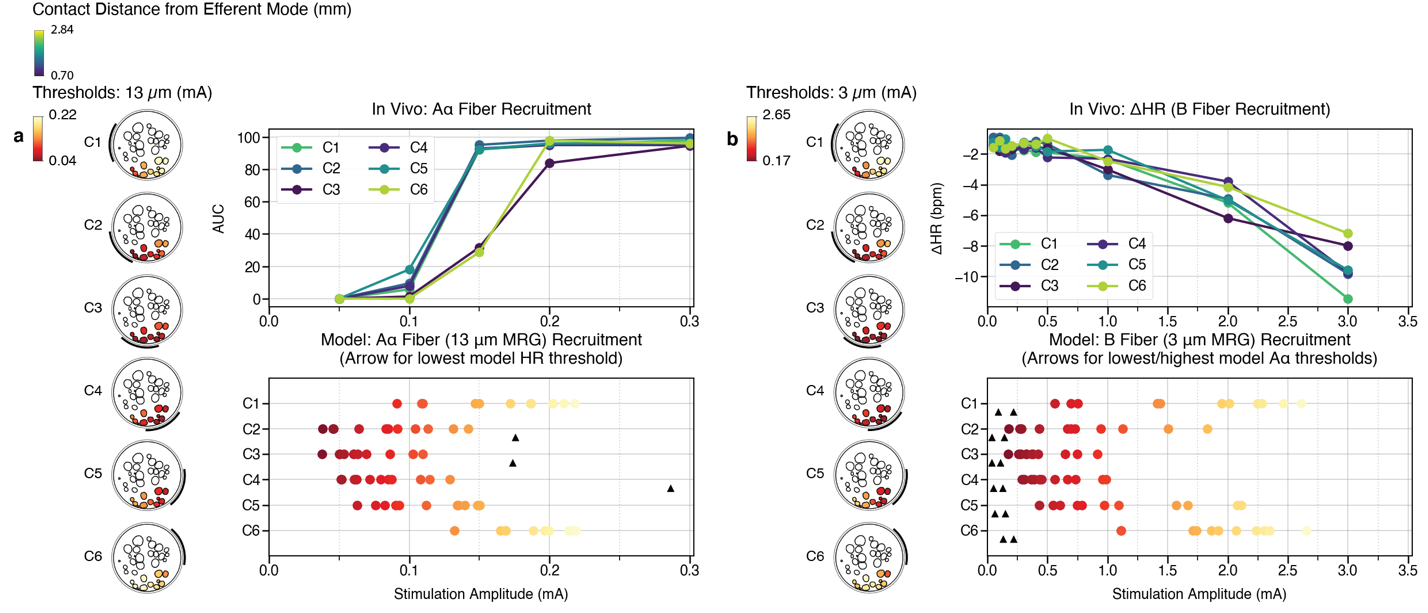

Figure 12. Comparison of in vivo thresholds to model thresholds in the pig cervical vagus nerve with the ImThera cuff electrode for Animal #3. a, Aα fiber thresholds modeled in the efferent mode compared to LIFE signals with corresponding conduction velocity. The arrows in the bottom panel indicate the lowest model B fiber threshold (i.e., heart rate onset threshold) for the corresponding contact, if the threshold fell on the x-axis range. In this case, all Aα fibers are activated before any model B fibers. b, B fibers modeled in the efferent mode compared to change in animal heart rate for a range of amplitudes. The arrows in the bottom panel indicate the lowest and highest model Aα threshold for the corresponding contact. In this case, all Aα fibers are activated before any model B fibers.

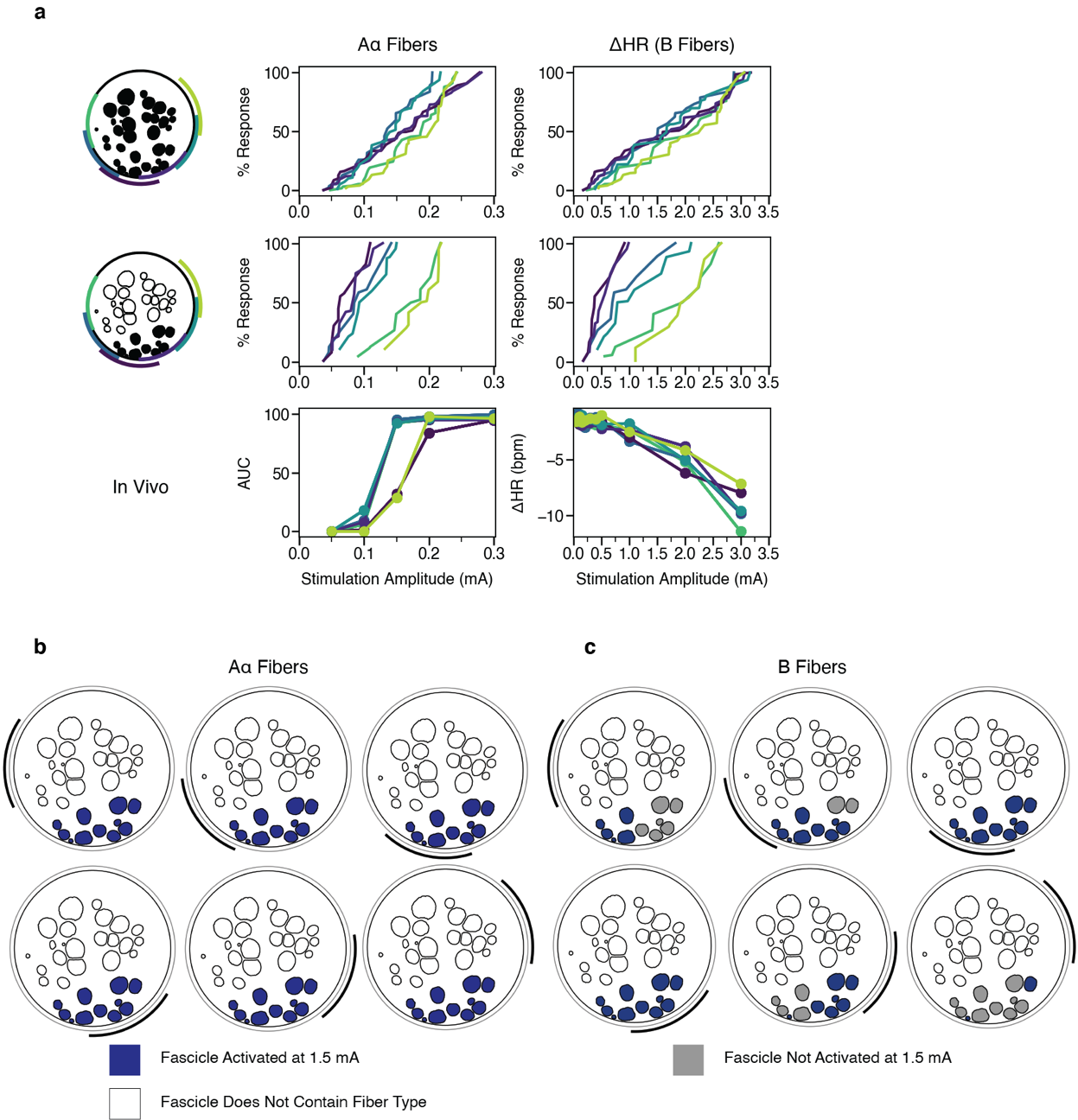

Figure 13. Animal #3 (with and without vagotopy) and in vivo recruitment curves for Aα and B fibers. a, Percent of modeled (top two rows) and in vivo (bottom row) Aα and B fibers activated versus stimulation amplitude across monopolar contact configurations (colors). In the top row of a, we modeled an Aα or B fiber in the centroid of each fascicle. In the middle row, we only modeled fibers in the black fascicles (i.e., the efferent mode). b-c, Fascicles labeled as suprathreshold (blue) and subthreshold (gray) in response to a 1.5 mA amplitude pulse for Aα and B fibers, respectively, for Animal #3. At this stimulation amplitude, Aα fibers are activated in all fascicles, while the B fibers are activated concomitantly only in the fascicles closest to the contact.

###### Animal #4

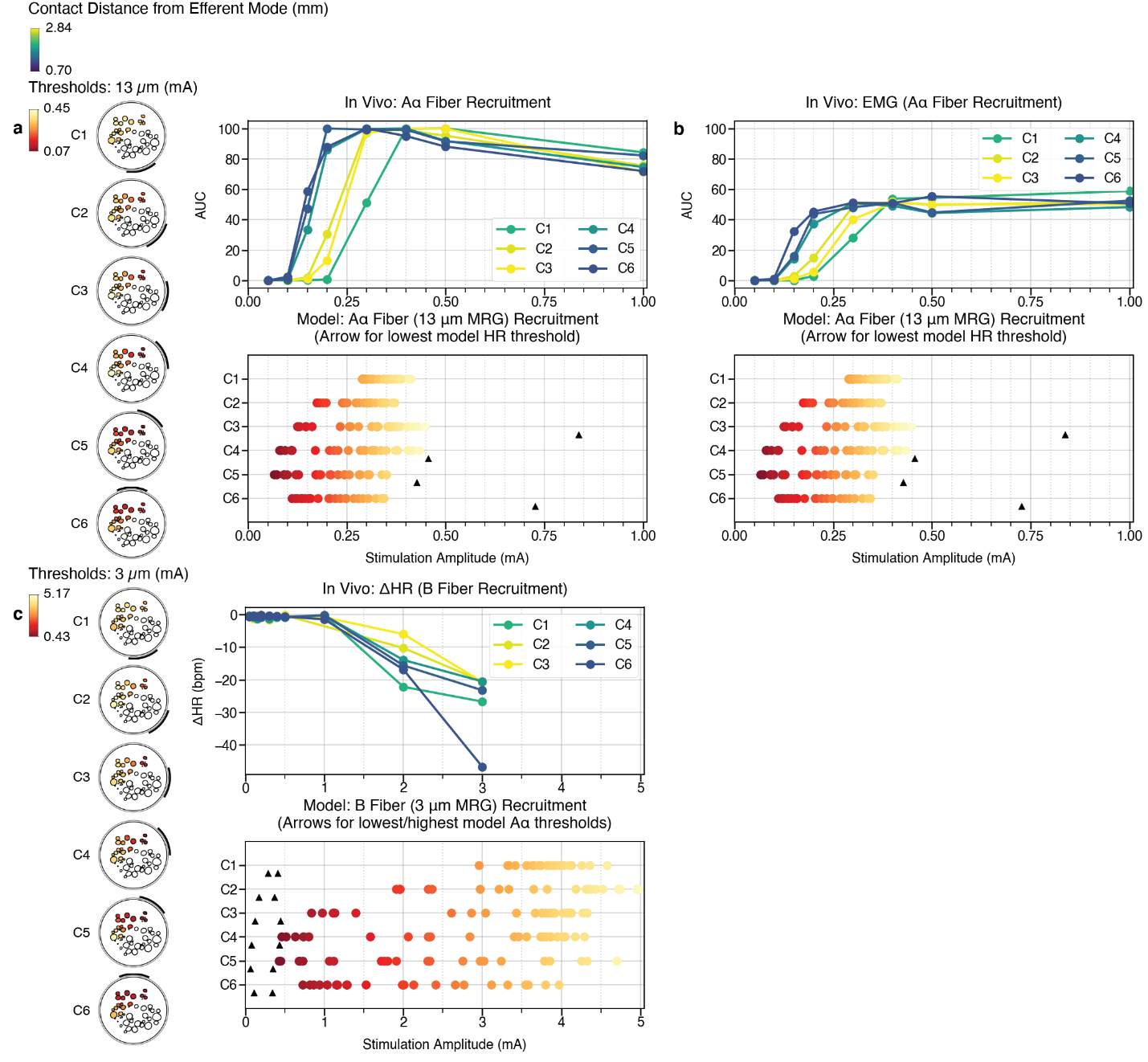

Figure 14. Comparison of in vivo thresholds to model thresholds in the pig cervical vagus nerve with the ImThera cuff electrode for Animal #4. a, Aα fiber thresholds modeled in the efferent mode compared to LIFE signals with corresponding conduction velocity. The arrows in the bottom panel indicate the lowest model B fiber threshold (i.e., heart rate onset threshold) for the corresponding contact, if the threshold fell on the x-axis range. In this case, all Aα fibers are activated before any model B fibers. b, Same modeling data as a plotted beside the laryngeal EMG signal. c, B fibers modeled in the efferent mode compared to change in animal heart rate for a range of amplitudes. The arrows in the bottom panel indicate the lowest and highest model Aα threshold for the corresponding contact. In this case, all Aα fibers are activated before any model B fibers.

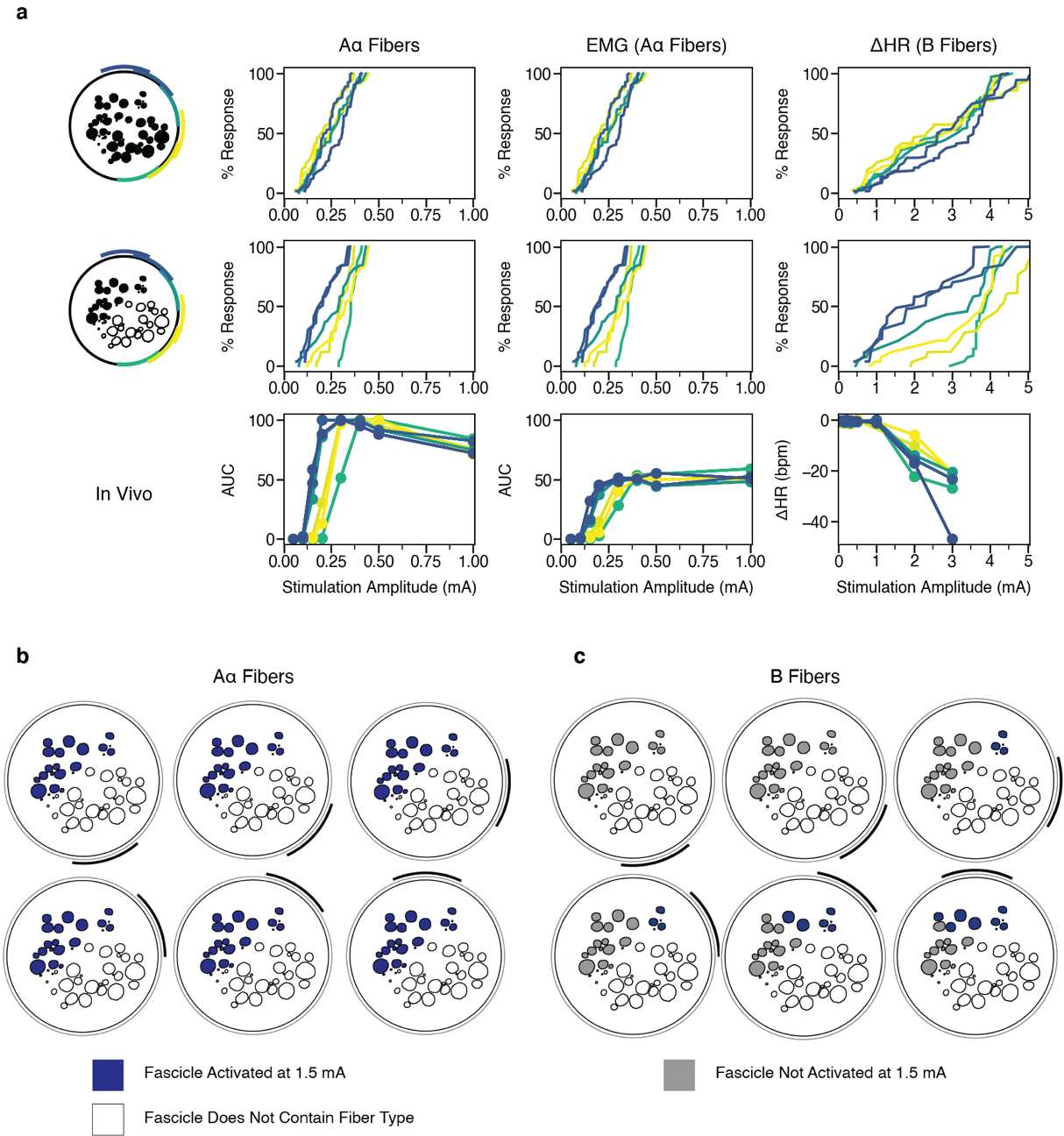

Figure 15. Animal #4 (with and without vagotopy) and in vivo recruitment curves for Aα and B fibers. a, Percent of modeled (top two rows) and in vivo (bottom row) Aα and B fibers activated versus stimulation amplitude across monopolar contact configurations (colors). In the top row, we modeled an Aα or B fiber in the centroid of each fascicle. In the middle row, we only modeled fibers in the black fascicles (i.e., the efferent mode). b-c, Fascicles labeled as suprathreshold (blue) and subthreshold (gray) in response to a 1.5 mA amplitude pulse for Aα and B fibers, respectively, for Animal #4. At this stimulation amplitude, Aα fibers are activated in all fascicles, while the B fibers are activated concomitantly only in the fascicles closest to the contact.

###### Animal #5

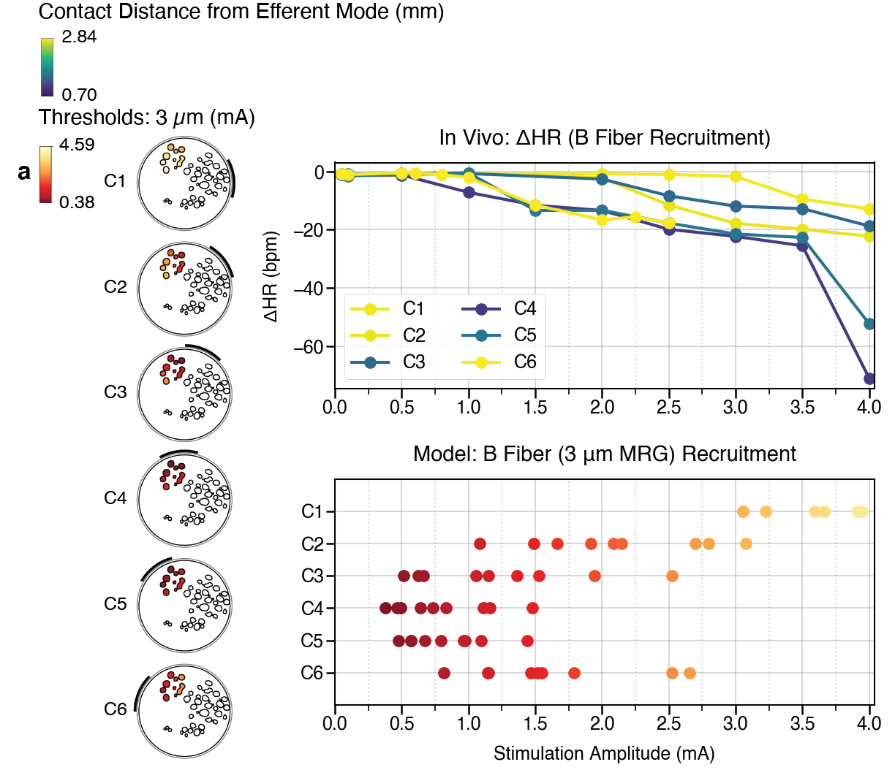

Figure 16. Comparison of in vivo thresholds to model thresholds in the pig cervical vagus nerve with the ImThera cuff electrode for Animal #5. a, B fibers modeled in the efferent mode compared to change in animal heart rate for a range of amplitudes.

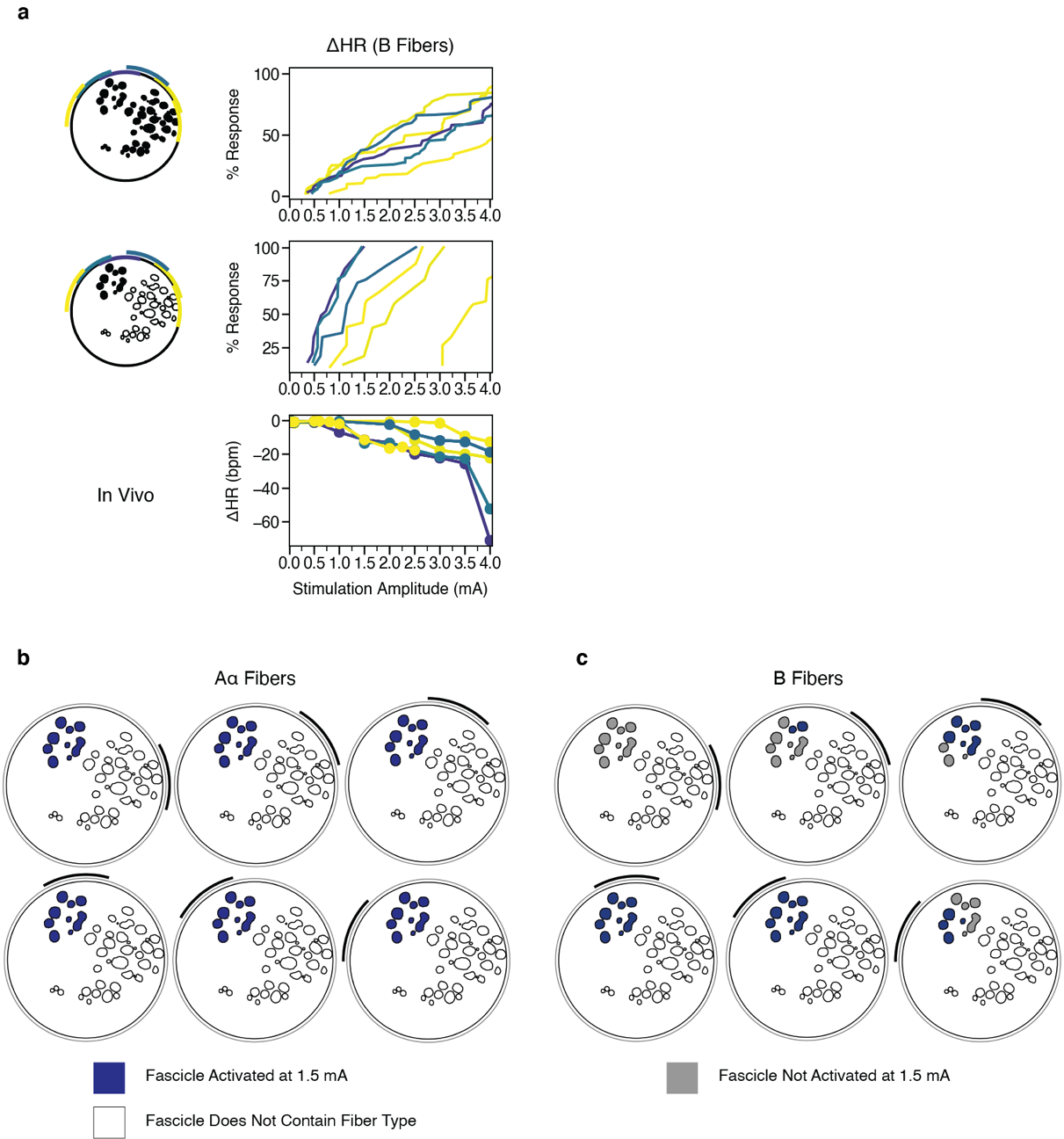

Figure 17. Animal #5 model and in vivo recruitment curves for B fibers. a, Percent of modeled (top two rows) and in vivo (bottom row) B fibers activated versus stimulation amplitude across monopolar contact configurations (colors). In the top row, we modeled a B fiber in the centroid of each fascicle. In the middle row, we only modeled fibers in the black fascicles (i.e., the efferent mode). b-c, Fascicles labeled as suprathreshold (blue) and subthreshold (gray) in response to a 1.5 mA amplitude pulse for Aα and B fibers, respectively, for Animal #5. At this stimulation amplitude, Aα fibers are activated in all fascicles, while the B fibers are activated concomitantly only in the fascicles closest to the contact.

###### Animal #6

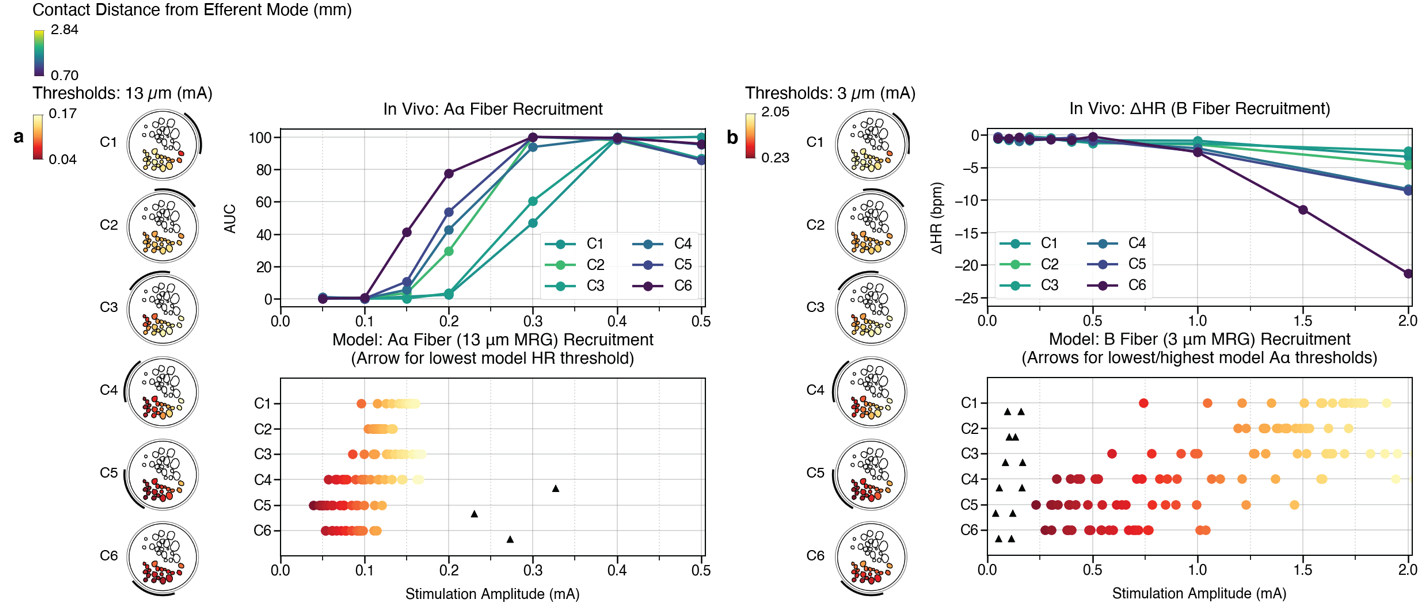

Figure 18. Comparison of in vivo thresholds to model thresholds in the pig cervical vagus nerve with the ImThera cuff electrode for Animal #6. a, Aα fiber thresholds modeled in the efferent mode compared to LIFE signals with corresponding conduction velocity. The arrows in the bottom panel indicate the lowest model B fiber threshold (i.e., heart rate onset threshold) for the corresponding contact, if the threshold fell on the x-axis range. In this case, all Aα fibers are activated before any model B fibers. b, B fibers modeled in the efferent mode compared to change in animal heart rate for a range of amplitudes. The arrows in the bottom panel indicate the lowest and highest model Aα threshold for the corresponding contact. In this case, all Aα fibers are activated before any model B fibers.

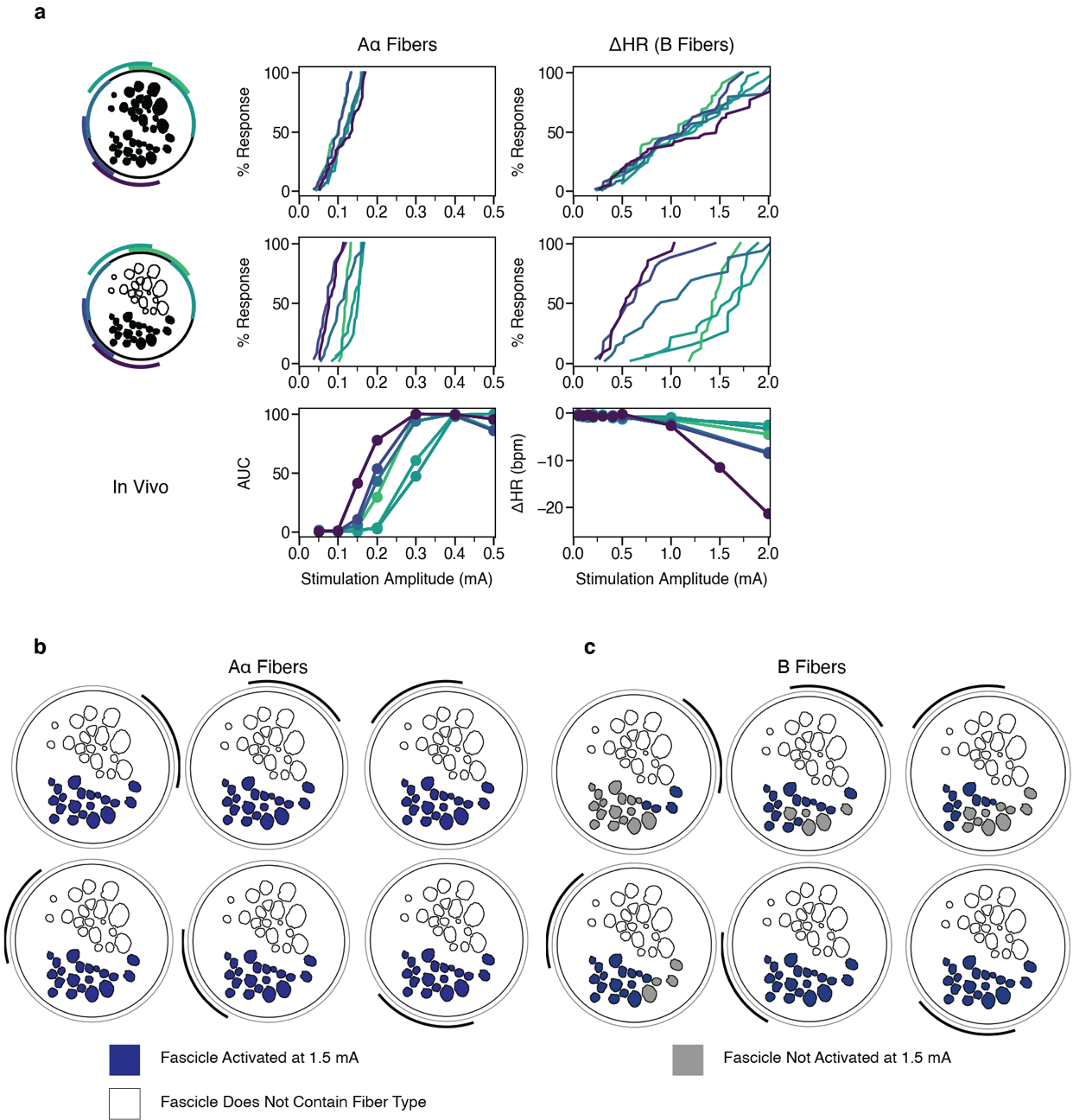

Figure 19. Animal #6 (with and without vagotopy) and in vivo recruitment curves for Aα and B fibers. a, Percent of modeled (top two rows) and in vivo (bottom row) Aα and B fibers activated versus stimulation amplitude across monopolar contact configurations (colors). In the top row of a, we modeled an Aα or B fiber in the centroid of each fascicle. In the middle row, we only modeled fibers in the black fascicles (i.e., the efferent mode). b-c, Fascicles labeled as suprathreshold (blue) and subthreshold (gray) in response to a 1.5 mA amplitude pulse for Aα and B fibers, respectively, for Animal #6. At this stimulation amplitude, Aα fibers are activated in all fascicles, while the B fibers are activated concomitantly only in the fascicles closest to the contact.

##### Off-Target Activation

The electrostatics generated by motor neurons may be recorded on various other recording electrodes, such as the LIFE implanted on the vagus (Figure 20). To ascertain the origin of the measured signal and account for the above-mentioned confounds, in a subset of animals (n=5), recordings were also taken from the CA and CT following the transection of the RL and following the transection of both the RL and SL (Figure 20, Figure 22). Transections of nerves innervating the CA and CT confirmed that activation of the motor fibers within these nerves was responsible for the elicited EMG response of the neck muscles. In general, transection of the recurrent laryngeal branch removed the slower component of EMG recordings, and SL transactions removed the faster component (Figure 20). Vagotomies were also performed to ensure that evoked compound action potentials recorded via LIFE were neural in source, as opposed to anticipated or unanticipated sources of artifact (Figure 20, Figure 22). A vagotomy between the stimulating and recording electrode clearly eliminated the recorded eCAP, definitively confirming the source as neural and not artifact (Figure 20).

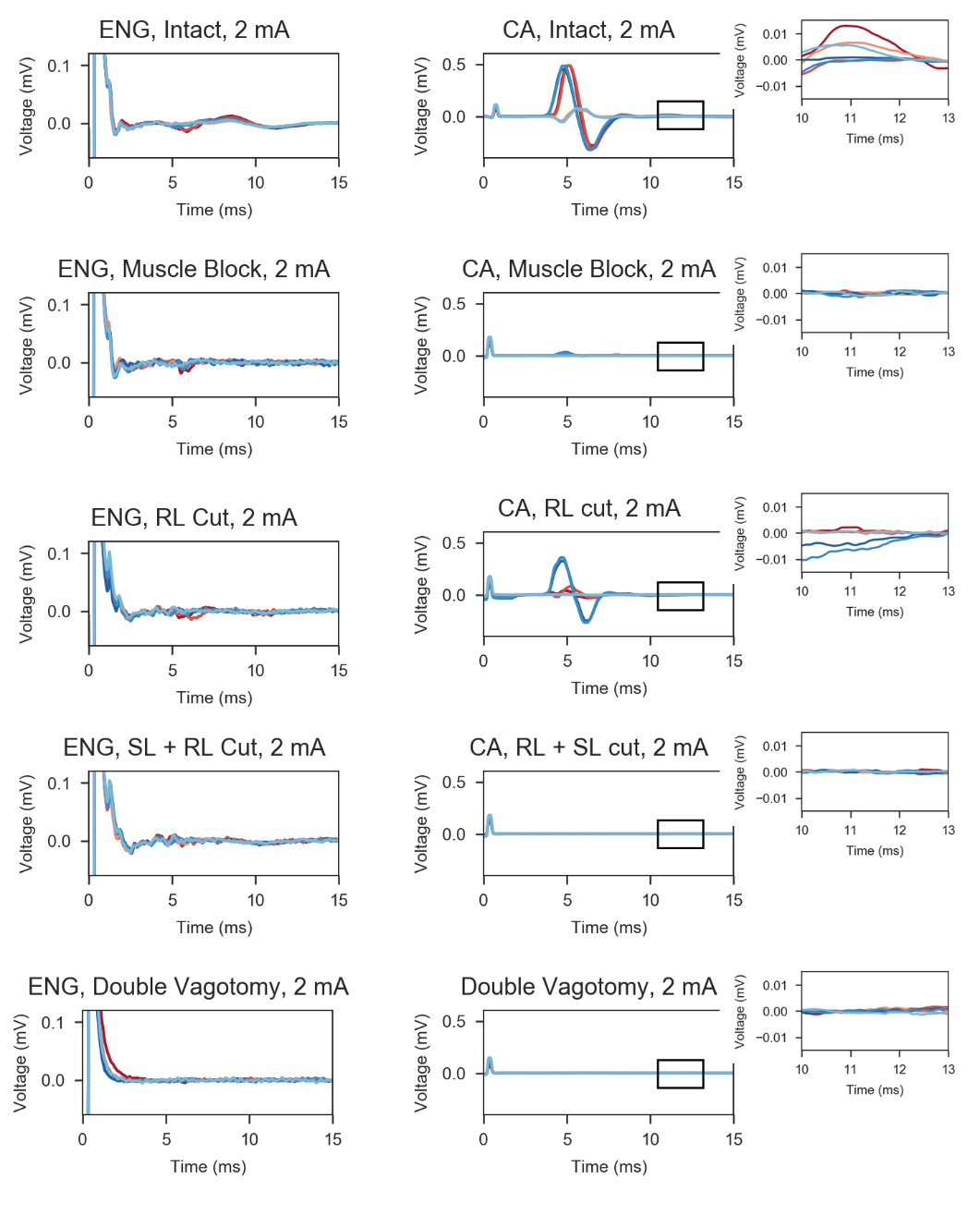

**Figure 20.** ENG vs EMG recordings from Animal #4. The left column shows the ENG from the vagus nerve. The right column shows the EMG from the cricoarytenoid (CA) muscle. Red/blue traces correspond to unique stimulation contacts. Each row illustrates an additional nerve transection. **Left.** Neural recordings show slight EMG artifact between 4 and 10 ms (top row), which disappears after transection of the SL and RL (second row), leaving exclusively neural signals. After the double vagotomy (bottom row), all neural traces disappear, as expected. **Right.** EMG recordings from the CA, showing innervation by both SL and RL (top row). Late EMG component (green arrows and zoomed panels) disappeared after RL transection (second row). No signs of EMG activity after both SL and RL were cut.

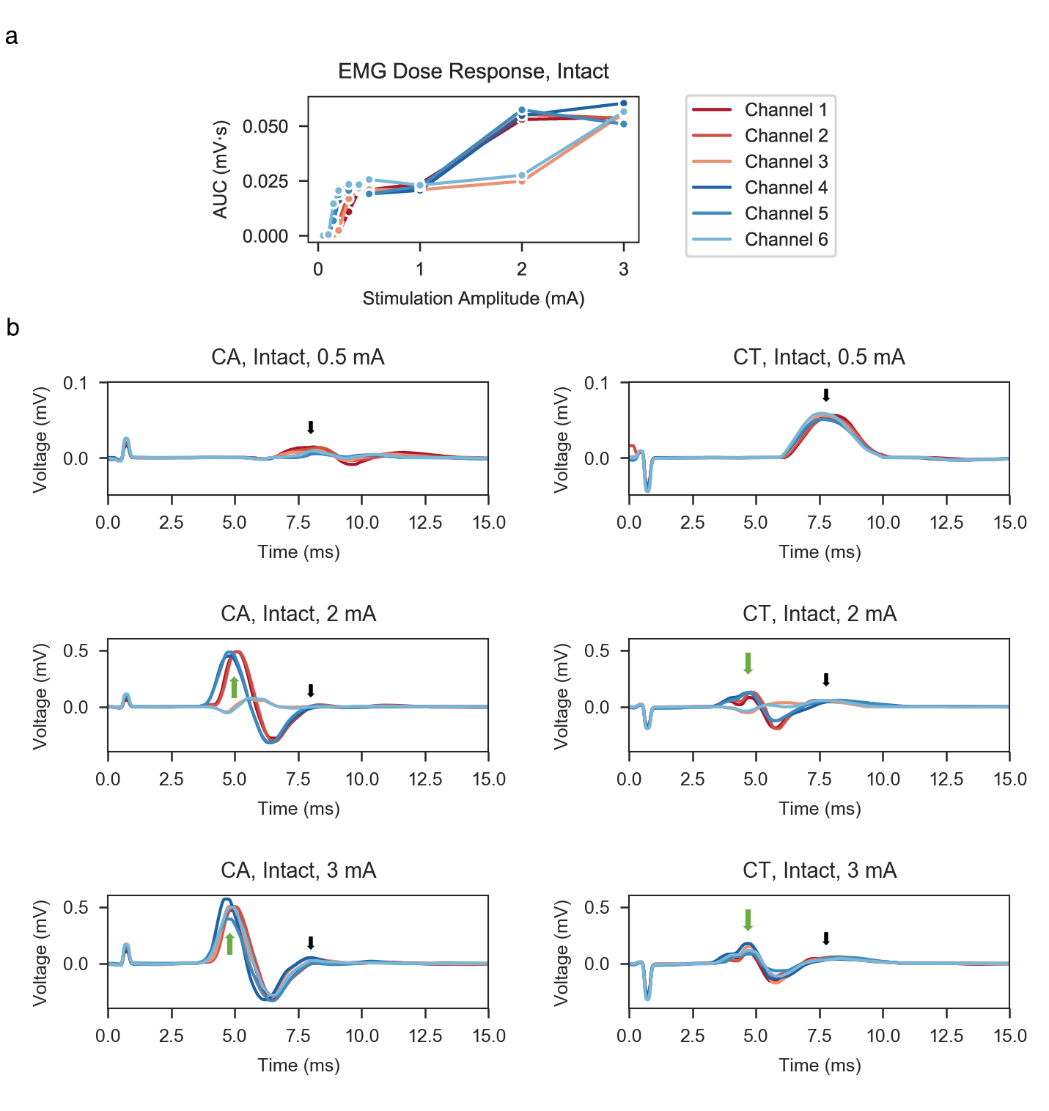

**Figure 21.** Dose-response curves and individual EMG traces from CA and CT muscles at three stimulation amplitudes from Animal #4. **A.** Dose-response curve for each stimulation contact, with two plateaus in the “Intact” condition: one below 1000 µA and one above 1 mA. The second plateau begins to saturate on select stimulating contact (1, 2, 4, and 5) around 2 mA and reaches full saturation on all channels by 3000 µA. **B.** EMG traces separated by columns according to muscle group (CA (left) and CT (right)) and by rows according to stimulation amplitude (0.5, 2, and 3 mA). Note the different y-axes below 2 mA stimulation. Two EMG components are present in the middle and bottom rows, characterized as ‘fast’ (green arrows) and ‘slow’ (black arrows) responses, due to activation of the SL fibers outside of the cuff and activation of the RL fibers within the cuff, respectively. Only the ‘slow’ component from direct innervation of the RL fibers is present in the top row. In particular, only select contacts produce indirect activation at 2 mA (1, 2, 4, and 5). All contact traces show full saturation of both components by 3 mA.

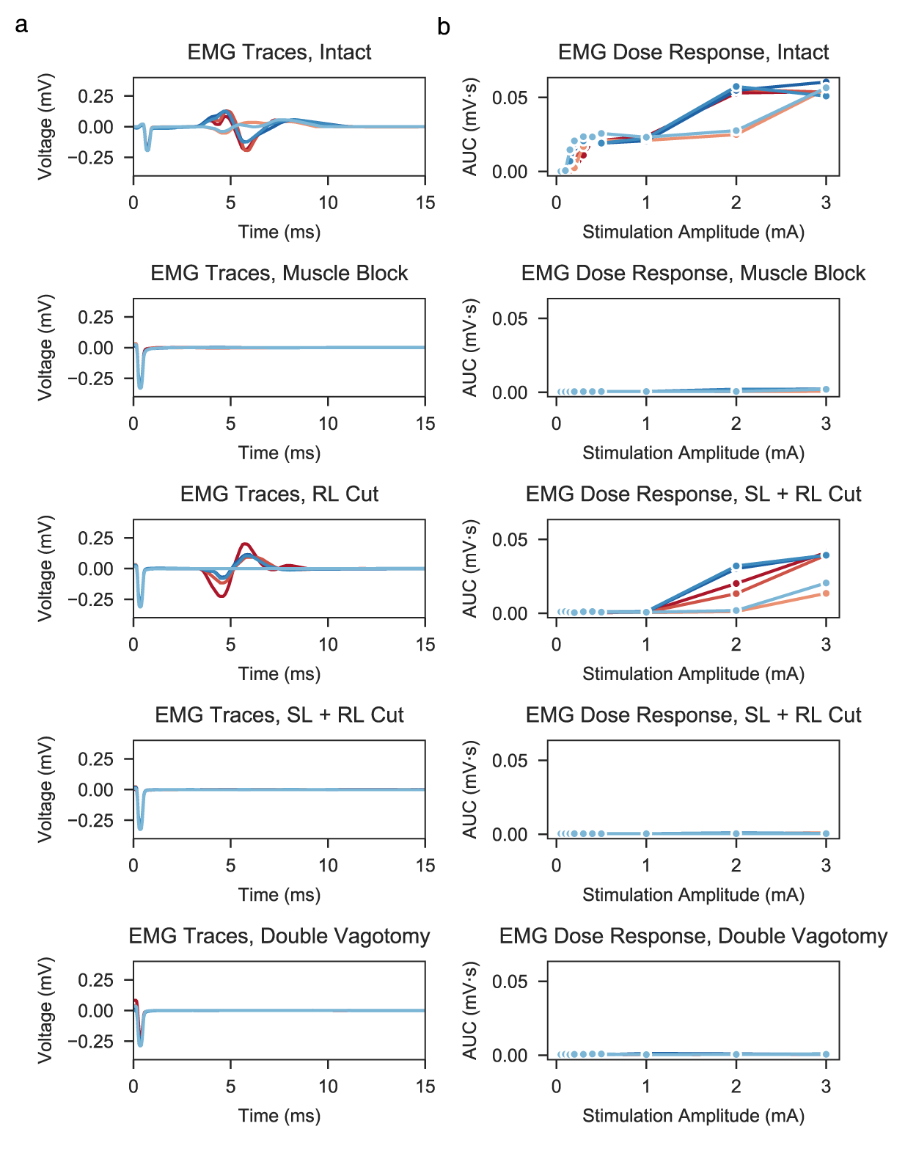

**Figure 22. a,** EMG responses from Animal #4 at the maximum stimulation amplitude delivered. Traces are presented in the intact nerve, muscle block, and SL + RL conditions to demonstrate effectiveness of muscle block and transections. **b,** EMG dose-response curves of conditions matching traces in the same row. Legend from figure 20 is used.

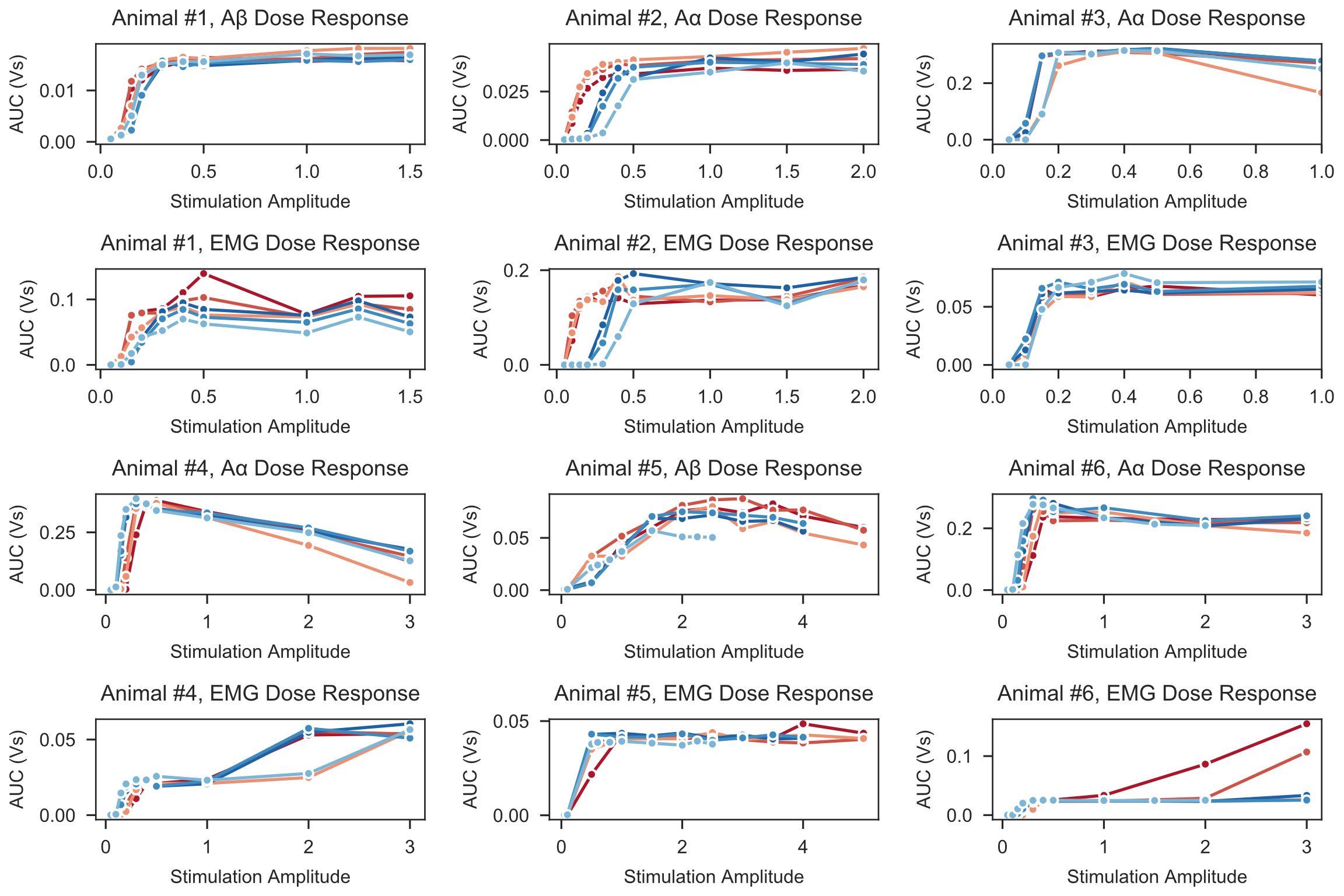

**Figure 23.** Paired ENG and EMG dose-response curves across the cohort. Aα AUCs were calculated in Animals # 2,3,4, and 6. In Animals #1 and #5, the stimulus artifact obscured the neural signal in this latency range; therefore, Aβ AUCs were calculated. Legend from Figure 20 is used.

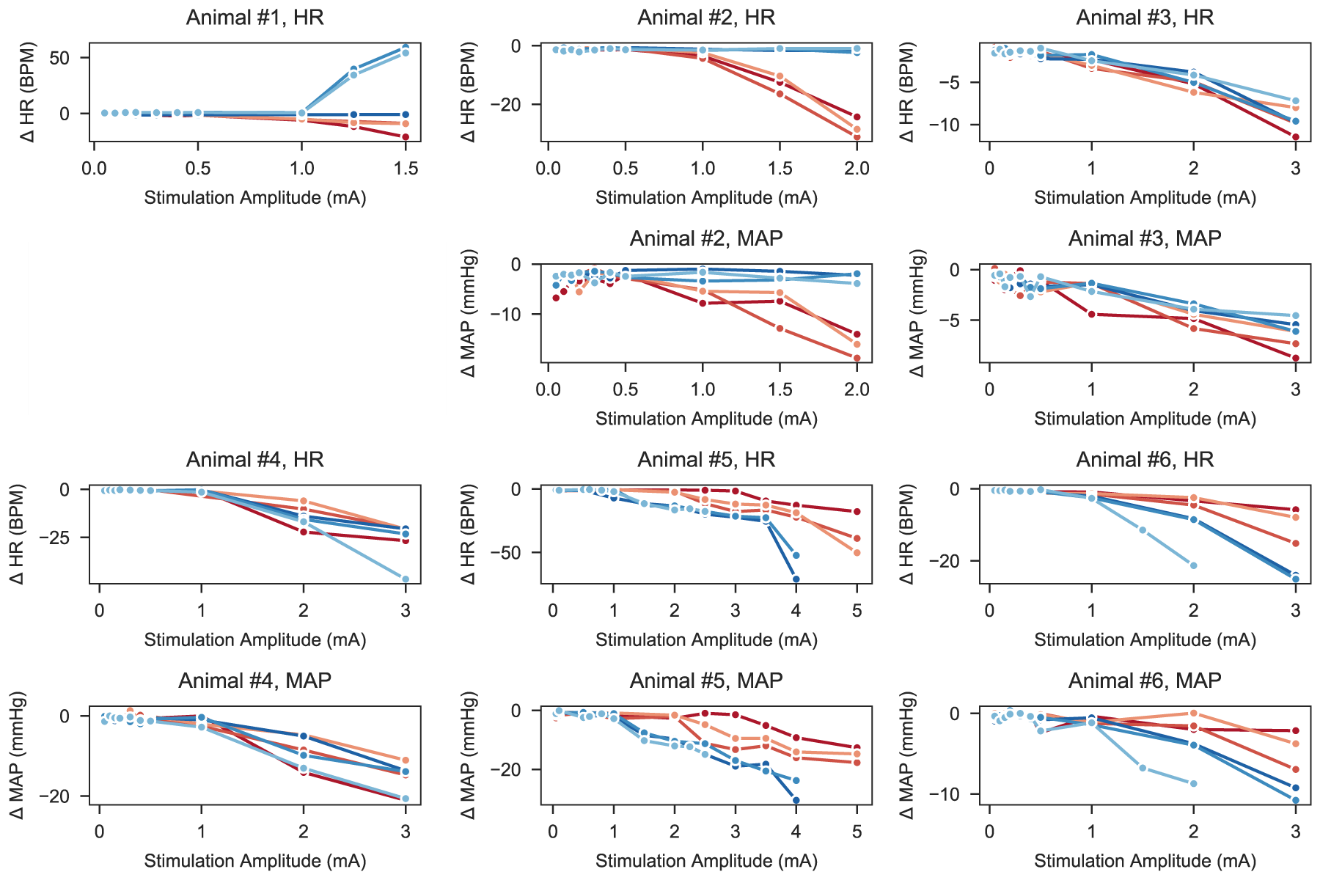

**Figure 25.** Dose-response curves comparing changes in heart rate (HR) and mean arterial pressure (MAP) across the cohort. The mean stimulation thresholds for HR and MAP responses are similar (1.312 ± 0.533 mA and 1.408 ± 0.724 mA, respectively). Red/blue traces correspond to unique stimulation contacts. Note the different x and y axes.

Boyd, I. A., & Kalu, K. U. (1979). Scaling factor relating conduction velocity and diameter for myelinated afferent nerve fibres in the cat hind limb. *The Journal of Physiology*, *289*, 277–297. https://doi.org/10.1113/jphysiol.1979.sp012737

Cogan, S. F. (2008). Neural Stimulation and Recording Electrodes. *Annual Review of Biomedical Engineering*, *10*(1), 275–309. https://doi.org/10.1146/annurev.bioeng.10.061807.160518

Fazan, V. P. S., & Lachat, J.-J. (1997). Qualitative and Quantitative Morphology of the Vagus Nerve in Experimental Chagas’ Disease in Rats: A Light Microscopy Study. *The American Journal of Tropical Medicine and Hygiene*, *57*(6), 672–677. https://doi.org/10.4269/ajtmh.1997.57.672

Hines, M. L., & Carnevale, N. T. (1997). The NEURON simulation environment. *Neural Computation*, *9*(6), 1179–1209. https://doi.org/10.1162/neco.1997.9.6.1179

Howell, B., & Grill, W. M. (2014). Evaluation of high-perimeter electrode designs for deep brain stimulation. *Journal of Neural Engineering*, *11*(4), 46026. https://doi.org/10.1088/1741-2560/11/4/046026

Licursi de Alcântara, A. C., Salgado, H. C., & Sassoli Fazan, V. P. (2008). Morphology and morphometry of the vagus nerve in male and female spontaneously hypertensive rats. *Brain Research*, *1197*, 170–180. https://doi.org/10.1016/j.brainres.2007.12.045

Manzano, G. M., Giuliano, L. M. P., & Nóbrega, J. A. M. (2008). A brief historical note on the classification of nerve fibers. *Arquivos de Neuro-Psiquiatria*, *66*(1), 117–119. https://doi.org/10.1590/S0004-282X2008000100033

McIntyre, C. C., Grill, W. M., Sherman, D. L., & Thakor, N. V. (2004). Cellular effects of deep brain stimulation: Model-based analysis of activation and inhibition. *Journal of Neurophysiology*, *91*(4), 1457–1469. https://doi.org/10.1152/jn.00989.2003

McIntyre, C. C., Richardson, A. G., & Grill, W. M. (2002). Modeling the excitability of mammalian nerve fibers: Influence of afterpotentials on the recovery cycle. *Journal of Neurophysiology*, *87*(2), 995–1006. https://doi.org/10.1152/jn.00353.2001

Mei, N., Condamin, M., & Boyer, A. (1980). The composition of the vagus nerve of the cat. *Cell and Tissue Research*, *209*(3), 423–431. https://doi.org/10.1007/BF00234756

Musselman, E. D., Cariello, J. E., Grill, W. M., & Pelot, N. A. (2021). ASCENT (Automated Simulations to Characterize Electrical Nerve Thresholds): A Pipeline for Sample-Specific Computational Modeling of Electrical Stimulation of Peripheral Nerves. *PLoS Computational Biology*.

Pelot, N. A., Behrend, C. E., & Grill, W. M. (2019). On the parameters used in finite element modeling of compound peripheral nerves. *Journal of Neural Engineering*, *16*(1), 16007. https://doi.org/10.1088/1741-2552/aaeb0c

Pelot, N. A., Goldhagen, G. B., Cariello, J. E., Musselman, E. D., Clissold, K. A., Ezzell, J. A., & Grill, W. M. (2020). Quantified Morphology of the Cervical and Subdiaphragmatic Vagus Nerves of Human, Pig, and Rat. *Frontiers in Neuroscience*, *14*, 1148. https://doi.org/10.3389/fnins.2020.601479

Petrossians, A., Whalen, J. J., Weiland, J. D., & Mansfeld, F. (2011). Surface modification of neural stimulating/recording electrodes with high surface area platinum-iridium alloy coatings. *2011 Annual International Conference of the IEEE Engineering in Medicine and Biology Society*, 3001–3004. https://doi.org/10.1109/IEMBS.2011.6090823

Ranck, J. B. (1975). Which elements are excited in electrical stimulation of mammalian central nervous system: A review. *Brain Research*, *98*(3), 417–440. https://doi.org/10.1016/0006-8993(75)90364-9

Rose, T. L., & Robblee, L. S. (1990). Electrical stimulation with Pt electrodes. VIII. Electrochemically safe charge injection limits with 0.2 ms pulses (neuronal application). *IEEE Transactions on Biomedical Engineering*, *37*(11), 1118–1120. https://doi.org/10.1109/10.61038

Soltanpour, N., & Santer, R. M. (1996). Preservation of the cervical vagus nerve in aged rats: Morphometric and enzyme histochemical evidence. *Journal of the Autonomic Nervous System*, *60*(1), 93–101. https://doi.org/10.1016/0165-1838(96)00038-0

Trevathan, J. K., Baumgart, I. W., Nicolai, E. N., Gosink, B. A., Asp, A. J., Settell, M. L., Polaconda, S. R., Malerick, K. D., Brodnick, S. K., Zeng, W., Knudsen, B. E., McConico, A. L., Sanger, Z., Lee, J. H., Aho, J. M., Suminski, A. J., Ross, E. K., Lujan, J. L., Weber, D. J., … Shoffstall, A. J. (2019). An Injectable Neural Stimulation Electrode Made from an In-Body Curing Polymer/Metal Composite. *Advanced Healthcare Materials*, *8*(23), 1900892. https://doi.org/10.1002/adhm.201900892

Van Rossum, G., & Drake, F. L. (2009). *Python 3 Reference Manual*. CreateSpace.

Weerasuriya, A., Spangler, R. A., Rapoport, S. I., & Taylor, R. E. (1984). AC impedance of the perineurium of the frog sciatic nerve. *Biophysical Journal*, *46*(2), 167–174. https://doi.org/10.1016/S0006-3495(84)84009-6

Wilks, S. J., Hara, S. A., Ross, E. K., Nicolai, E. N., Pignato, P. A., Cates, A. W., & Ludwig, K. A. (2017). Non-clinical and Pre-clinical Testing to Demonstrate Safety of the Barostim Neo Electrode for Activation of Carotid Baroreceptors in Chronic Human Implants. *Frontiers in Neuroscience*, *11*. https://doi.org/10.3389/fnins.2017.00438
